# Positive selection in gamete interaction proteins in *Carnivora*

**DOI:** 10.1101/2022.03.22.485370

**Authors:** Francisco Pisciottano, Clara Campos, Clementina Penna, Carlos David Bruque, Toni Gabaldón, Patricia Saragüeta

## Abstract

The absence of robust interspecific isolation barriers among pantherines, including the iconic South American jaguar (*Panthera onca*), led us to study molecular evolution of typically rapidly evolving reproductive proteins within this subfamily and related groups. In this study, we delved into the evolutionary forces acting on the Zona Pellucida (ZP) gamete interaction protein family and the sperm-oocyte fusion protein pair IZUMO1-JUNO across the *Carnivora* order, distinguishing between *Caniformia* and *Feliformia* suborders and anticipating few significant diversifying changes in the *Pantherinae* subfamily. A chromosome-resolved jaguar genome assembly facilitated coding sequences, enabling the reconstruction of protein evolutionary histories. Examining sequence variability across more than 30 Carnivora species revealed that *Feliformia* exhibited significantly lower diversity compared to its sister taxa, *Caniformia*. Molecular evolution analyses of ZP2 and ZP3, subunits directly involved in sperm-recognition, unveiled diversifying positive selection in *Feliformia*, *Caniformia*, and *Pantherinae*, although no significant changes were linked to sperm binding. Structural cross-linking ZP subunits, ZP4, and ZP1 exhibited lower levels or complete absence of positive selection. Notably, the fusion protein IZUMO1 displayed prominent positive selection signatures and sites in basal lineages of both *Caniformia* and *Feliformia*, extending along the *Caniformia* subtree but absent in *Pantherinae*. Conversely, JUNO did not exhibit any positive selection signatures across tested lineages and clades. Eight Caniformia-specific positive selected sites in IZUMO1 were detected within two JUNO-interaction clusters. Our findings provide for the first time insights into the evolutionary trajectories of ZP proteins and the IZUMO1-JUNO gamete interaction pair within the *Carnivora* order.

## 1. Introduction

The zona pellucida (ZP) is the extracellular matrix synthesized in the ovaries of mammals during follicular development. It is found surrounding the oocyte and early embryo plasma membrane and constitutes a three-dimensional structure composed of ZP glycoprotein filaments. Oocyte formation, fertilization and early embryo development are biological processes involving ZP proteins (Yanagimachi, 1994; Dean, 2007; Wassarman & Litscher, 2009; Gupta and Bhandari, 2011; Gupta et al., 2012; Tanihara et al., 2013; Shu et al., 2015; Moros-Nicolás et al., 2018a; Moros-Nicolás et al., 2021). These proteins are representative components involved in reproduction and are usually found evolving under positive selection. It is proposed that this favoured rapid evolution plays an important role in reproductive isolation of diverging taxa (Swanson et al., 2001; Swanson & Vacquier, 2002; Turner & Hoekstra, 2008, Meslin et al., 2012).

ZP matrix of most, but not all, eutherians is composed of four ZP protein subunits: ZP1-4 (Figure 1A). ZP3 was the first glycoprotein of the ZP to be considered indispensable for fertilization in mammals (Bleil & Wassarman, 1980). Independent research findings revealed that mouse oocytes lacking a functional *Zp3* gene exhibit infertility and the absence of the ZP (Liu et al., 1996; Rankin et al., 1996). As a result, ZP3 garnered broad consensus in the scientific community as the predominant and indispensable protein responsible for sperm binding. However, further research revealed that ZP2 is also a protein essential for mouse fertility as it is required for sperm binding (Rankin et al., 2001, Avela et al., 2014). Capacitated sperm attaches to an N-terminal domain of ZP2 in the extracellular ZP surrounding eggs (Baibakov et al., 2012; Avella et al., 2014). After fertilization, ovastacin, an oocyte-specific metalloendoproteases, is exocytosed from egg cortical granules and cleaves extracellular ZP2 (Burkart et al., 2012). The cleavage site for this event is located in the ZP-N2 domain of ZP2. Post-fertilization proteolytic cleavage at the N terminus of ZP2 protein prevents sperm binding to the ZP and provides a block to polyspermy (Baibakov et al., 2012). ZP1 is a ZP protein not directly related to gamete interaction, but its function is no less relevant for the formation of the ZP. ZP1 protein is involved in ZP protein filaments cross-linking and interacts with other ZP proteins, particularly ZP2 and ZP3, to form the ZP matrix, giving and maintaining its structure (Monné & Jovine, 2011; Wassarman & Litscher, 2018; Nishimura et al., 2019; Stsiapanava et al., 2020). Therefore, the main function of ZP1 protein is to contribute to the structure and organization of the ZP. ZP4 was the last of the eutherian ZP proteins to be identified (Lefièvre et al., 2004), as it is not present in the house mouse (*Mus musculus*), the most extended mammalian model in which most studies of the ZP had been performed. ZP4 is considered a paralog of ZP1 and the origin of these two modern genes can be traced back to a duplication of a common ancestral gene in the early evolution of amniotes (Goudet et al., 2008). Like ZP1, ZP4 plays a structural and mechanical role that is not essential during fertilization *in vivo* but fundamental for the protection of the developing embryo (Lamas-Toranzo et al., 2019).

**Figure 1:**
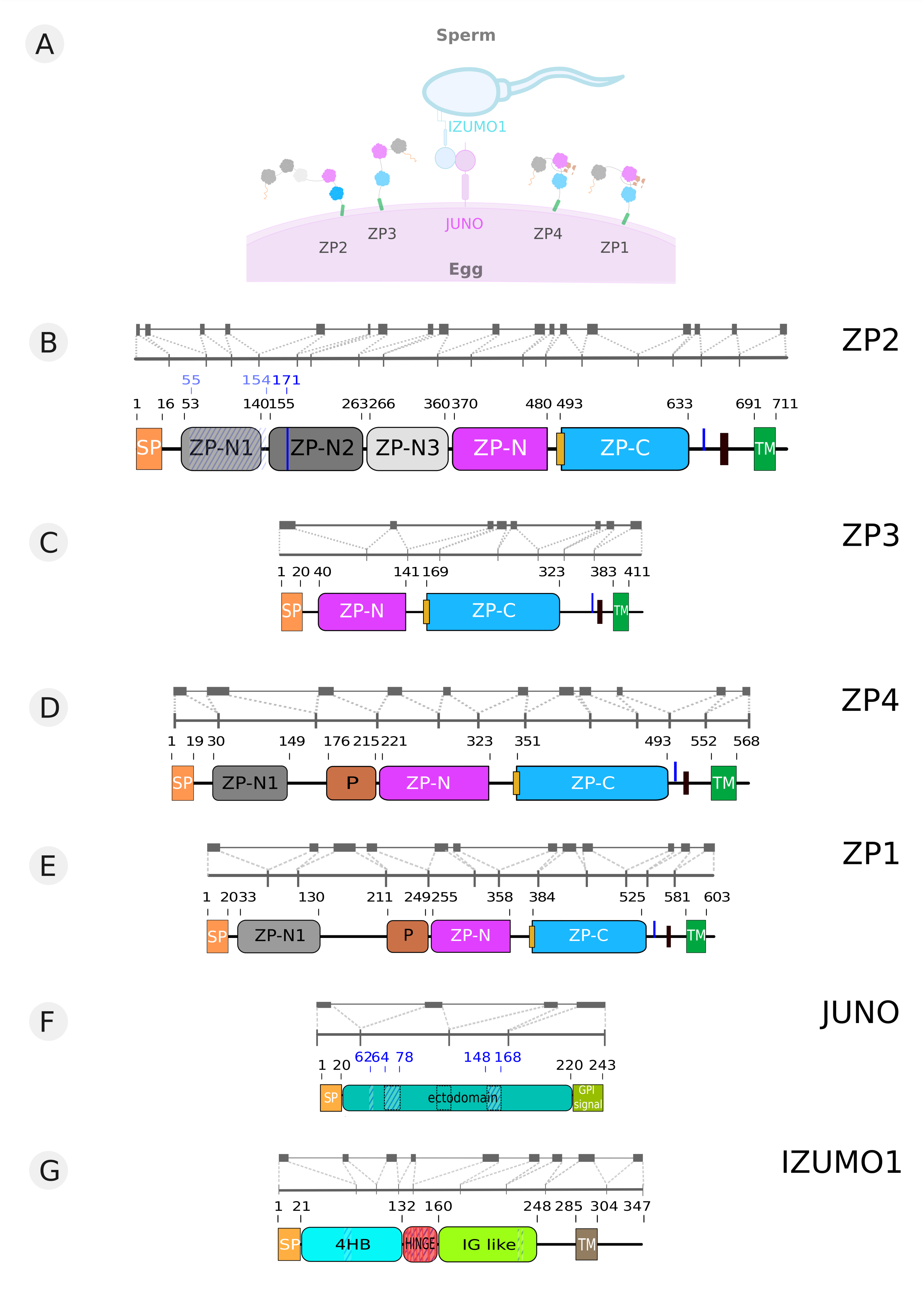
Schematic overview of ZP and JUNO-IZUMO1 gamete interaction proteins. A. Diagram of sperm and egg showing studied gamete interaction proteins: zona pellucida (ZP) proteins ZP1 to 4, JUNO from oocyte and IZUMO1 from sperm. B. Schematic representation of jaguar *Zp2* gene, illustrating the arrangement of exons and introns (first line). Visualization of the mature transcript of the *Zp2* gene (second line). Schematic representation of the protein domains comprising ZP2 with jaguar start and end positions for each domain (third line). Start and end positions of the putative sperm-binding region are indicated with blue numbering above the protein scheme. SP: Signal Peptide; ZP-N1/2/3: ZP-N like domains; ZP-N and ZP-C: Zona Pellucida N-terminal and C-terminal domains; TM: Transmembrane domain. Blue vertical full line in ZP-N2: ZP2 ovastatin cleavage site. Blue vertical line after ZP-C: furin cleavage site. Diagonal striped blue pattern: ZP2 sperm-binding region. Yellow rectangle: internal hydrophobic patch. Black rectangle: external hydrophobic patch. C. Schematic representation of jaguar *Zp3* gene, illustrating the arrangement of exons and introns (first line). Visualization of the mature transcript of the *Zp3* gene (second line). Schematic representation of the protein domains comprising ZP3 with jaguar start and end positions for each domain (third line). SP, ZP-N, ZP-C, TM, blue vertical line after ZP-C, yellow and black rectangles, idem B. D. Schematic representation of jaguar *Zp4* gene, illustrating the arrangement of exons and introns (first line). Visualization of the mature transcript of the *Zp4* gene (second line). Schematic representation of the protein domains comprising ZP4 with jaguar start and end positions for each domain (third line). P: Trefoil domain. SP, ZP-N1, ZP-N, ZP-C, TM, blue vertical line after ZP-C, yellow and black rectangles, idem B. E. Schematic representation of the *Zp1* gene, illustrating the arrangement of exons and introns (first line). Visualization of the mature transcript of the *Zp1* gene (second line). Schematic representation of the protein domains comprising ZP1 with jaguar start and end positions for each domain (third line). P: Trefoil domain. SP, ZP-N1, ZP-N, ZP-C, TM, blue vertical line after ZP-C, yellow and black rectangles, idem B. F. Schematic representation of jaguar *Juno* gene, illustrating the arrangement of exons and introns (first line). Visualization of the mature transcript of the *Juno* gene (second line). Schematic representation of the protein domains comprising JUNO with jaguar start and end positions for each domain and start and end positions of the regions involved in interaction with IZUMO1 in blue numbers (third line). Flexible regions are demarcated on the domains with a black dashed rectangles. Regions involved in interaction with the IZUMO1 are shown with blue stripes diagonal pattern. SP: Signal Peptide; GPI signal: signal for anchor addition. G. Schematic representation of jaguar *Izumo1* gene, illustrating the arrangement of exons and introns (first line). Visualization of the mature transcript of the *Izumo1* gene (second line). Schematic representation of the protein domains comprising IZUMO1 with jaguar start and end positions for each domain (third line). Regions involved in interaction with the JUNO are shown with blue stripes diagonal pattern. SP: Signal Peptide; HB: N-terminal alpha helices bundle; HINGE: Hinge domain; IG like: C-terminal immunoglobulin-like domain; TM: Transmembrane domain.

ZP2-ZP3 filaments are formed through heterodimerization of the ZP2 and ZP3 proteins after their secretion to the extracellular space by the oocyte. These heterodimers then assemble end-to-end, forming long filaments that extend throughout the ZP matrix (Jovine et al., 2002; Wassarman & Litscher, 2018). ZP1 forms a scaffold for the assembly of the ZP2-ZP3 filaments, aiding in their arrangement and providing structural integrity to the ZP matrix. ZP4 is a minor component of the ZP matrix but is thought to have a role in mediating intermolecular interactions between ZP2-ZP3 filaments. ZP4 may act as a link between adjacent ZP2-ZP3 filaments, contributing to the overall stability and architecture of the ZP matrix (Wassarman & Litscher, 2018).

The structure of ZP proteins is characterized by different regions and domains crucial for the secretion and assembly of ZP filaments. These are an N-terminal secretory signal peptide (SP) that identifies secreted proteins; the ZP module, a conserved sequence of ~260 amino acids composed of a ZP-N and a ZP-C domain, and include 8 or 10 invariant cysteine residues that form four, or alternatively five, disulphide bonds; a recognition site for members of the proprotein convertase family of proteolytic enzymes, a consensus furin cleavage site (CFCS); and a C-terminal propeptide that includes a single-spanning transmembrane (TM) domain (Jovine et al., 2005; Monné et al., 2006; Bokhove et al., 2016; Wilburn & Swanson, 2017). ZP3 protein presents the simplest domain composition among all members of the ZP protein family as it displays the basic structural elements common to all ZP proteins: a signal peptide, one complete ZP module, divided into its N-terminal and C-terminal moieties, a furin cleavage site and a hydrophobic C-terminal trans-membrane domain (TMD), all of them necessary for their secretion and incorporation in the ZP matrix. In contrast to the minimalistic domain composition of ZP3, the ZP2 protein stands out for having the most extensive domain count among all members of the ZP protein family. Alongside the distinctive ZP module found in all ZP proteins, which consists of N-terminal and C-terminal domains, ZP2 possesses three extra ZP-N domains situated at the N-terminal region preceding the complete ZP module. These additional domains go from ZP-N1 to ZP-N3. Beyond the ZP protein common structural organization of domains, ZP1 displays a single ZP-N domain and a trefoil domain, which is important for the cross-linking between ZP filaments (Greve and Wassarman, 1985; Nishimura et al., 2019). Given that ZP4 and ZP1 are paralogous genes resulting from the duplication of a common ancestral gene, both keep the same domain structure and have a 45 aa long trefoil domain and a ZP-N1 subdomain in their amino-terminal region (Goudet et al., 2008).

The evolutionary history of the ZP protein family reveals numerous gene gains and losses. During the early evolution of amniotes, the duplication event giving rise to *Zp1* and *Zp4* occurred (Hughes and Barratt, 1999; Bausek et al., 2000; Goudet et al., 2008). Subsequent pseudogenization events led to modern placental mammals, which possess a ZP structure composed of three subunits instead of four. Pseudogenization events are more prevalent in *Zp1* gene compared to other ZP genes across placental mammal species. *Zp1* loss has been observed in certain species of the *Caniformia* suborder, including the dog (*Canis familiaris*), dingo (*Canis lupus dingo*), and fox (*Vulpes vulpes*) (Goudet et al., 2008; Moros-Nicolás et al., 2018a). Pseudogenization of *Zp1* gene also occurred in various Pinniped species such as the Antarctic fur seal (*Arctocephalus gazella*) and the Weddell seal (*Leptonychotes weddellii*) (Moros-Nicolás et al., 2018a). Additional instances of *Zp1* pseudogenization are found in the evolution of Cetartiodactyla, where many even-toed ungulates and cetaceans lack a functional version of this gene (Goudet et al., 2008; Stetson et al., 2012). Furthermore, pseudogenization events affecting *Zp1* have been reported in certain lineages of small primates, such as marmosets and tarsiers (Stetson et al., 2012). In contrast, the pseudogenization of *Zp4* gene has been documented in the house mouse (*Mus musculus*) and other members of the Mus genus (Lefièvre et al., 2004; Izquierdo-Rico et al., 2021). However, it is important to note that ZP3 and ZP2 proteins have not been reported as lacking in any mammalian species, indicating their essential function in gamete interaction and fertilization (Liu et al., 1996; Rankin et al., 1996, 2001).

Considerable efforts have been made to elucidate the sperm interacting partners of egg ZP proteins. However, no sperm surface proteins have been confirmed to interact with egg ZP proteins and be indispensable for fertilization. One of the most well known cases is the ZP3 receptor (ZP3R) protein. ZP3R was initially proposed to be essential for mouse fertilization. However, knockout studies in mice have shown that ZP3R is not indispensable for the fertilization process, suggesting either its non-involvement in sperm-ZP binding or the presence of functionally redundant proteins sperm-egg interaction (Muro et al., 2012). To date, only one essential mammalian sperm-egg interaction protein pair has been identified: IZUMO1 on sperm interacts with egg membrane-anchored JUNO to facilitate binding (Inoue et al., 2005; Bianchi et al., 2014) (Figure 1A). IZUMO1 was first identified through a systematic proteomic analysis of proteins enriched in the sperm membrane (Inoue et al., 2005). Knockout studies in mice have highlighted the indispensability of IZUMO1 in mediating sperm-egg fusion (Inoue et al., 2005). Structural analyses have identified the IZUMO domain within IZUMO1 as a key component for binding to the oocyte-specific glycoprotein JUNO and initiating the fusion process (Aydin et al., 2016; Ohto et al., 2016). Given its significant role in fertilization, the protein was aptly named “IZUMO1” after a Japanese shrine associated with marriage, emphasizing its critical function in facilitating sperm-egg interaction and highlighting its symbolic relevance to the deity Izumo, who is traditionally believed to preside over marriages and relationships (Inoue et al., 2005). JUNO protein, also known as folate receptor 4 (FOLR4), is encoded by the gene *Izumo1r*. Originally discovered in 2014 and renamed JUNO after the Roman queen goddess of marriage and childbirth, it is a transmembrane glycoprotein that serves as the counterpart in the oocyte to the sperm protein IZUMO1. The interaction between JUNO and IZUMO1 is essential for sperm to dock to the surface of an egg (Bianchi et al., 2014), a crucial recognition step prior to membrane fusion (Inoue et al., 2015). Although neither protein has been conclusively shown to be a fusogen, there is recent evidence that IZUMO1 may have fusogenic activity (Brukman et al., 2023).

The selected non-model species of study - the jaguar (*Panthera onca*)-is the most important predator in the South American biome and in danger of extinction in Argentina (Paviolo et al., 2016). Hence, it is crucial to develop genomic, evolutionary, and functional studies about this species. Genomic introgression analysis carried out together with the first release of the jaguar complete genome exposed a complex pattern of historical inter-specific hybridization in the *Pantherinae* lineage (Figueiró et al., 2017). This and other precedents, such as the existence in captivity of natural hybrids between extant pantherines (Yadav et al., 2019) and the high efficiency in obtaining IVF hybrids of oocytes with whole Zona Pellucida between different felines (Duque Rodriguez et al., 2021), are consistent with relaxed inter-specific isolation barriers in this group. The lack of emergence of isolation mechanisms between *Pantherinae* species would have allowed the historical process of introgression among them.

The disruption of interaction or fusion between gametes due to changes in the counterpart proteins is one of many mechanisms through which prezygotic isolation can arise. If such changes were to occur in these proteins and were fixed by reinforcement mechanisms, they would be detected in the gene sequence encoding these proteins as episodic positive selection signatures. In other words, there would be an increase in the fixation rate of non-synonymous changes compared to synonymous changes in a particular lineage, followed by the successive conservation of these changes. Beside, increased variability at a key protein site essential for interaction with its counterpart would be detected as diversifying positive selection in the clade and could also lead to prezygotic isolation. While these are not the only reasons why reproductive proteins may exhibit signatures of positive selection, if such changes occurred, we would expect them to be located in functionally important regions.

Traits of coevolution between sperm and egg protein counterparts were reported to conduct reproductive isolation (Swanson et al., 2003; Aagaard et al., 2006). Up till now, IZUMO1-JUNO is the only gamete interaction protein pair that has been identified to be essential for fertilization in mammals. This work focuses on ZP and IZUMO1-JUNO proteins evolution in order to look into their role in speciation in the pantherines. For this purpose, *Feliformia* suborder was taken as the phylogenetic background of the pantherines and compared to its sister clade, *Caniformia*. Therefore, the phylogenetic context of this work is the whole *Carnivora* order, with special attention to *Caniformia*-*Feliformia* differences and to the evaluation of selection forces acting on the *Pantherinae* subfamily. Hence, we hypothesize that sperm-oocyte interaction and fusion proteins are particularly conserved or at most accumulate few diversifying changes across *Pantherinae*, in accordance with the weak or absent isolating mechanisms in this lineage.

## 2. Materials and Methods

### 2.1 Sequence retrieval and alignment

All *Carnivora* species coding sequences of the studied gamete-interacting proteins were downloaded from Ensembl (Howe et al., 2021) and GenBank Nucleotide database (Supplementary tables S2-S7). The number of species from caniforms and feliforms was balanced for all analyses. Therefore, HiC chromosome-length DNA Zoo genomes of Black footed cat (*Felis* nigripes), Jaguarundi (*Herpailurus yagouaroundi*), Clouded leopard (*Neofelis nebulosa*), Snow leopard (*Panthera* uncia), Fishing cat (*Prionailurus viverrinus*) and Leopard cat (*Prionailurus bengalensis*) were downloaded and sequences were obtained via local BLAST searches and manually assembled. Jaguar (*Panthera onca*) sequences were obtained from the first version of the jaguar reference genome (Figueiró et al, 2017) and the HiC assembled jaguar genome (DNA zoo) by performing a local BLASTn using leopard (*Panthera pardus*) ortholog sequence as query, obtained from GenBank (NCBI Resource Coordinators, 2016). After a manual inspection and identification of sequencing or assembly errors of both genomes, the sequences that exhibited less fragmentation and missing data were selected. Visual inspection and multiple sequence alignments were performed with MEGA 5.2 (Kumar et al., 2018) using Clustal and/or Muscle (Li, 2003; Edgar, 2004). A careful manual curation of the alignments was performed in order to maximize quality of aligned sequences. Low quality sequences that could not be replaced or significantly improved were discarded. Low quality or non-aligning portions of coding sequences in one species were deleted and, whenever possible, replaced using additional sequence sources such as Nscan and GeneScan, accessed through UCSC Genome Browser in order to guarantee homology and the quality of the aligned sequences.

### 2.2 Positive selection analysis

In order to detect signatures of molecular adaptation the modified Model A branch-site test 2 of positive selection (Zhang et al., 2005) was used. This model is available in codeml program from the PAML4 package (Yang, 2007). This model has 2 variants: the focal branch variant, used to detect episodic positive selection, and the subtree variant, used to detect divergent evolution. In each case this model is able to detect signatures of positive selection in preassigned specific branches or clades respectively, and in both cases these changes can be assigned to individual codon positions. Both models were applied for each of the tested groups in this work: *Feliformia*, *Caniformia* and *Pantherinae.* By appropriately labeling the basal lineage of each group, we were able to assess the presence of episodic selection specifically at the basal lineage of the group or the presence of diversifying selection across the entire clade derived from that basal lineage.

Test 2 of the Model A of codeml consists of two nested hypotheses that must be compared: in the alternative hypothesis, positive selection is allowed only in the selected branch or clade, but not in the background branches or clades; and in the null hypothesis no positive selection is allowed in any branch or clade of the tree. The heterogeneity among protein sites was handled by site classes with different omega values (ω=dN/dS). In this model three omega parameters were used: ω_0_<1, ω_1_=1 (fixed) and ω_2_>1. In the alternative hypotheses site classes with ω_2_ were enabled in the foreground branch or clade allowing the presence of positive selection, but in the null hypotheses only site classes with ω_0_ and ω_1_ were enabled, restricting sites to purifying selection or neutral evolution only. The fit of the gene data (sequence alignment and tree) to these hypotheses was obtained by maximum likelihood (ML) and evaluated by a likelihood ratio test (LRT) in which a statistic calculated from twice the difference of the ML obtained was compared to a chi-square distribution (1 degree of freedom) to infer the significance between hypotheses.

All positive selection analyses were performed using multiPAML 2.5. An in-house script developed to efficiently and easily run a large number of PAML analyses testing for multiple hypotheses in one or many genes in a single script run. The multiPAML 2.5 software code is publicly available at https://github.com/fpisciottano.

### 2.3 Gene trees reconstruction

Gene trees were reconstructed by ML from the sequence alignment data using the Jones-Taylor-Thornton (JTT) amino acid substitution model with partial deletion treatment for missing data. Although tree topologies were defined in these ways, branch lengths of the trees used in the branch-site positive selection analyses were estimated using M0 (one ratio) model from PAML package. Given that this test for positive selection detection is a very parameter-rich model, it was implemented by following an ad-hoc methodology designed to prevent local minimum solutions and maximize rerun results consistency. Instead of running all parameters optimization while evaluating the selection model, we pre-computed all branch lengths under the Goldman & Yang codon-based evolutionary model (Goldman & Yang, 1994) using M0 model, also known as one-ratio model, from PAML package. Thus the total number of parameters was split and only model-specific parameters were optimized during the positive selection analysis.

### 2.4 Positive selected sites

Positive selected sites were obtained by the Bayesian posterior analysis (BEB) implemented in codeml’s branch-site specific positive selection test *aposteri* of the maximum likelihood estimates (Yang et al., 2005). As site-specific models are sensitive to codon parameters, we repeated each analysis 6 times using 2 different codon equilibrium frequencies estimation parameter variants (CodonFreq parameter in PAML). Sites reported correspond to those consistent in all 3 replicates in each of the 2 CodonFreq variants applied (‘codon table’ and ‘3×4’) with a p-value of at least 0.8 in all reruns.

### 2.5 Sequence Identities

Mean sequence identities for the taxonomic groups of interest were obtained from pairwise sequence identities comparisons from each of the alignments using Sequence Identity And Similarity online tool (SIAS – http://imed.med.ucm.es/Tools/sias.html). Caniforms vs feliforms mean identity comparison for each protein was performed through a two sample T-Test (Welch’s T-test). Violin plot graphs for identity figures were drawn with Python using the Seaborn library (version 0.9.1). Statistical relevance of mean sequence identities differences between proteins was assessed by ANOVA and Scheffé post-hoc multiple comparisons were conducted.

### 2.6 Predicted structure model of ZP proteins

The predicted structural models for jaguar ZP proteins were performed with the DeepMind AlphaFold2 (Jumper et al., 2021) as implemented in ColabFold (Mirdita *et al*., 2021) using 20 templates (phenix versión). The University of California in San Francisco (UCSF) Chimera v1.16 program (Pettersen et al, 2004) was used for visualization and the backbone-dependent rotamer library was used for structural interpretation (Shapovalov et al., 2011). N-glycosylation and O-glycosylation sites were obtained from the UniProt database (The UniProt Consortium, 2021) and converted to jaguar positions through the multiple sequence alignments. The Coulombic Surface Colouring calculates electrostatic potential according to Coulomb’s law (coulombic surface colouring, −10 to +10 kcal/(mol*e), red to blue).

### 2.7 Protein domain identification

Protein domains and their corresponding positions in jaguar were defined using specific associated bibliography (Nishimura et al., 2019; Raj et al., 2017; Han et al., 2010; Moros-Nicolás et al., 2018a; Aydin et al., 2016; Kato et al., 2016) and our highly curated sequence alignments for position conversion along ortholog sequences. Start and end positions of ZP protein domains were later corrected based on the three-dimensional models and the observed and expected secondary structure of each domain. Secondary structure prediction for each ZP protein was obtained using PDBsum web server (Laskowski et al., 1997), PSIPRED 4.0 (Jones, 1999) and DISOPRED3 (Jones et al., 2014) and contrasted with the canonical ZP hemi-domains topology (Bokhove et al., 2018) in order to check if obtained domains fulfilled the structural characteristic features in number and type of secondary structures and also if they displayed all disulphide-bonds described as necessary for each domain correct folding. Lineal schematic representation showing ZP protein domains were based on Bokhove et al., 2018.

## 3. Results

The study of protein evolution has greatly changed thanks to the growing availability of complete genome sequences. For the jaguar, our no-model system of study, global protein evolution analysis is now possible owing to the availability of the jaguar genome draft (Figueiró et al., 2017). Furthermore, we have complemented and improved our evolutionary analyses of gamete interaction proteins using a second version of jaguar chromosome-resolved genome assembled by Hi-C long-range sequencing (Dudchenko et al., 2017).

The analyses start with an extensive compilation of all accessible coding sequences for the orthologs of gamete interaction proteins ZPs and IZUMO1-JUNO in all *Carnivora* species for which sequencing data is available in biological databases (Supplementary tables S2-S7). Jaguar sequences for ZP proteins and the IZUMO1-JUNO pair were manually assembled and reported for the first time in this paper (see online supplementary data). The complete sequences of the jaguar *Zp2*, *Zp3*, *Zp4 and Zp1* genes were obtained using BLASTn searches over the jaguar chromosome-resolution Hi-C genome assembly and/or the Illumina sequenced first draft of the jaguar genome assembly (Figueiró et al., 2017) (Supplementary table S1).

### 3.1 Orthologous sequences retrieval and multiple sequence alignment construction

*Panthera pardus* (leopard) orthologous coding sequences were used as queries in each case (*Zp2* GeneID: 109255525, *Zp3* GeneID: 109250568, *Zp4* GeneID:109248980, *Zp1* GeneID:109246518). Jaguar *Zp2* coding sequence comprises 2,151 nucleotides distributed across 18 coding exons (Figure 1B), exhibiting a conserved exon arrangement among all sequenced felid species, except for the jaguarundi (*Puma yagouaroundi*) ortholog, which displays 19 exons, and two puma (*Puma concolor*) alternative transcripts (XM_025916776.1 and XM_025916778.1), both containing an additional exon. For the jaguar *Zp3* gene, attempts to obtain the sequence from the first version of the jaguar genome were unsuccessful, yielding only a partial and fragmented sequence. However, using the complete sequence derived from the *Panthera onca* Hi-C genome assembly, it was confirmed that the jaguar *Zp3* gene is composed of 1,272 nucleotides distributed across eight exons (Figure 1C), consistent with all pantherine species orthologs and also with most sequenced felid species, except for the puma (*Puma concolor*) ortholog, which contains an additional exon, totalling nine exons. As for the jaguar *Zp4* gene coding sequence, it consists of 1,707 nucleotides organized into 11 exons (Figure 1D), encoding a 569 amino acid-long polypeptide. The complete coding sequence for Jaguar *Zp4* was obtained by conducting a BLASTn search using the leopard *Zp4* sequence as the query. Finally, the jaguar *Zp1* gene encompasses 1,818 nucleotides across 12 exons (Figure 1E), mirroring the exon quantity, grouping, and distribution pattern observed in other sequenced felids. Regarding JUNO protein coding gene, *Izumo1r,* complete sequence was obtained through local BLASTn over the Illumina genome assembly (Figueiró et al., 2017) and the chromosome-resolution Hi-C genome assembly using leopard ortholog coding sequence as query (GeneID:109267358). Jaguar *Izumo1r* gene is only 729 nucleotides in length and is composed of 4 coding exons (Figure 1F) and its locus structure, number and disposition of exons is well conserved along all available felid species. Jaguar JUNO protein is 243aa long peptide that starts with a 19aa long signal peptide and contains a folate receptor domain between its positions 26 and 201. On the other hand, the jaguar coding sequence of *Izumo1*, it was obtained through a local BLASTn of the lion *Izumo1* sequence (Ensembl gene ID: ENSPLOG00000009340) over the Hi-C genome assembly of *Panthera onca*. Jaguar *Izumo1* coding sequence is composed of 1,037 nucleotides distributed in 9 exons (Figure 1G) and encodes a 346 aa long protein.

In addition to individual jaguar sequences of the target genes, we carried out a comprehensive search, collection, assembly, and curation of all ortholog sequences available for these genes in *Carnivora* species (Supplementary tables S2-S7). It is worth noting that none of the sequences used in this study have been reported to meet high-quality standards or undergone manual annotation to date. Consequently, all sequences required rigorous review and curation during the multiple sequence alignment building process. As a consequence of the approach chosen to compensate caniforms and feliforms number of species (see Materials and Methods section), the multiple sequence alignments (MSA) achieved displayed a more balanced representation between caniforms and feliforms (see online supplementary data). The majority of the compiled final alignments comprise 31 species, with a roughly equal distribution of caniform and feliform sequences: 15 caniforms and 16 feliforms for *Zp2*, 16 caniforms and 15 feliforms for *Zp3*, 15 caniforms and 16 feliforms for *Zp4* and 15 caniforms and 16 feliforms for *Izumo1r*.. Among the feliform sequences, these alignments include six pantherine species, encompassing all five species within the *Panthera* genus as well as the clouded leopard of the *Neofelis* genus. However, in the *Zp1* alignment, only 28 species are present due to pseudogenization of sequences from species within the *Canidae* family, allowing the use of only 12 caniform and 16 feliform species The *Izumo1* alignment is the other exception, resulting in a total of 33 sequences: 17 caniforms and 16 feliforms. These sequences were later used for a sequence diversity characterization of the studied proteins through identity analysis. We contrasted the levels of molecular variability of these proteins within the order *Carnivora*. Next, we further examined the two suborders -*Caniformia* and *Feliformia*-that constitute the *Carnivora* order. These MSAs were also the basis for phylogeny reconstructions and the positive selection analyses reported further below.

### 3.2 Gamete Interaction Protein Identity

To begin the evolutionary study of ZP proteins and the IZUMO1-JUNO pair, a general characterization of their evolution rates was determined, exploring the variability within the *Caniformia* order. The sequence identity revealed that the ZP proteins exhibited comparable levels of variability, except for ZP1, which displayed significant differences from ZP2 and ZP3, but not from ZP4. Similarly, the JUNO protein showed variability levels comparable to those of the ZP proteins. In contrast, IZUMO1 displayed a notably higher variability when compared to all other studied proteins, presenting significant differences from all of them (Figure 2A and Supplementary table S8). Additionally, the levels of variability among groups within the order *Carnivora* for each protein showed that feliform sequences were similar to one another, while caniform sequences exhibited a more pronounced variability. The highest degree of similarity was observed in the sequences of the *Pantherinae* subfamily. Welch’s T-test over the mean identity values of the amino acid variability indicated that all the studied proteins exhibited significant differences between caniforms and feliforms (Figure 2B and Supplementary table S9). The statistical difference in ZP4 identity values between caniforms and feliforms (Welch’s T-test p-value = 0.03649) proved to be smaller in comparison to the differences observed in the other proteins (Welch’s T-test p-values: ZP2: 1.26e^-4^; ZP3: 2.57e^-4^; ZP1: 4.25e^-4^; IZUMO1: 1.28e^-4^; JUNO: 8.1e^-5^) (Supplementary table S9). No significant differences were detected between the mean identities of caniforms and feliforms for the control ACTN1 protein shown at the bottom of Figure 2B (Supplementary table S9).

**Figure 2:**
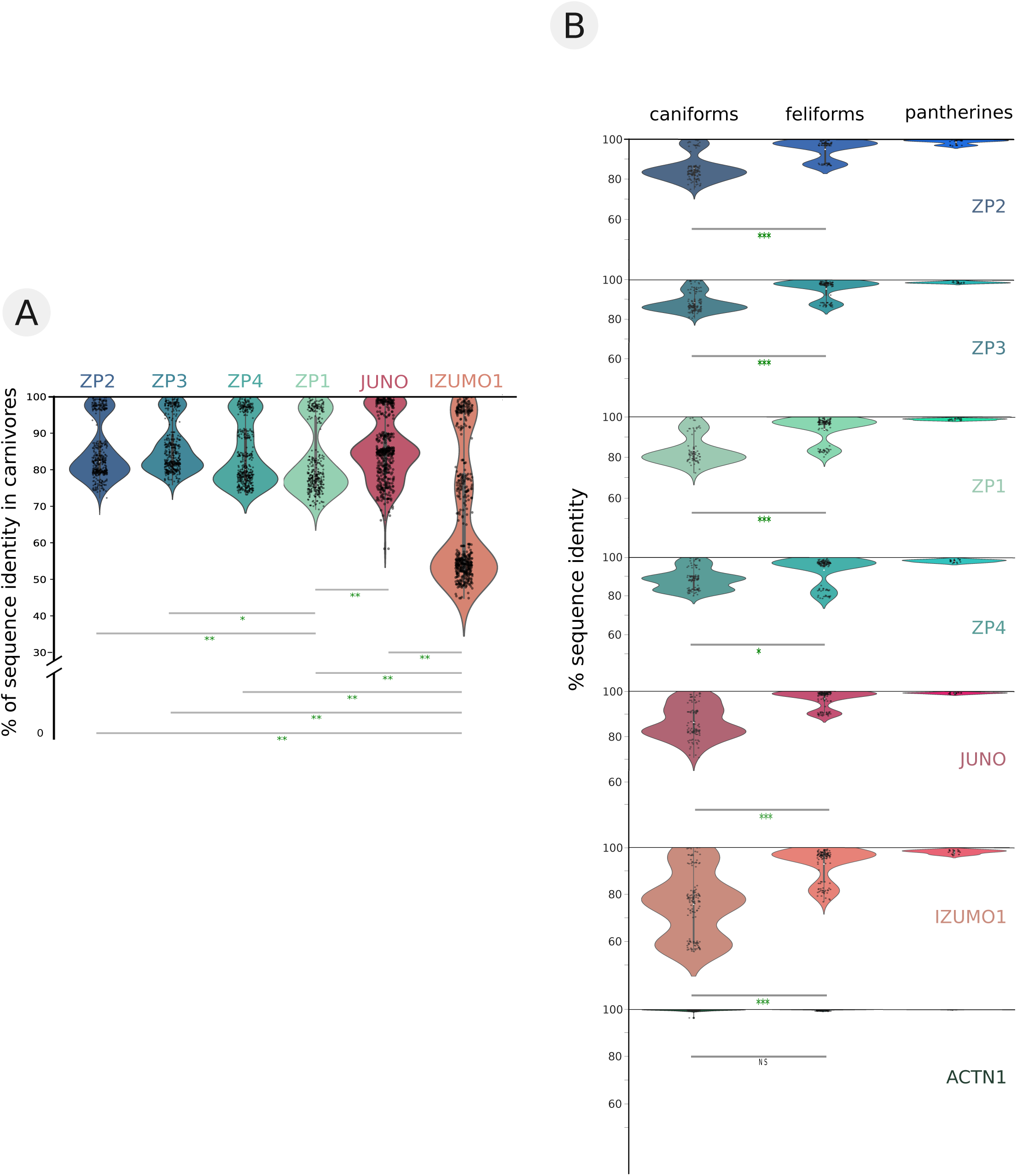
Identity profiling of ZP and JUNO-IZUMO1 gamete interaction proteins. A. ZP proteins, JUNO and IZUMO1 carnivores sequence identities. Violin plot showing pairwise sequence identities percentages (black dots) for ZPs, JUNO, and IZUMO1 proteins. Mean percentage of sequence identities for each protein is represented in a white dot. Scheffé’s T-test significance: (*) p<0.05 (**); p<0.01. B. ZPs, JUNO and IZUMO1 sequence identities for taxonomic groups of interest (caniforms, feliforms and pantherines). Violin plot showing pairwise sequence identities percentages (black dots). Mean percentage of sequence identity for each group is represented in a white dot. Welch’s T-test significance: (*) p<0.05 (**); p<0.01.

The positive selection analyses began with ZP isoforms directly involved in sperm-recognition, ZP2 and ZP3, and continued with the analysis of structural cross-linker ZP proteins, ZP4 and ZP1. Finally, we extended the analyses to the sperm–egg fusion pair IZUMO1-JUNO. Two control proteins: one under non-selection and another under previously reported positive selection in jaguar were added.

### 3.3 ZP proteins Positive Selection Analyses

The positive selection analyses revealed diversifying positive selection effects on the primary receptors ZP2 and ZP3 within the suborders of *Carnivora*. Specifically, ZP2 exhibits positive selection in the *Feliformia* suborder, while ZP3 shows positive selection in the *Caniformia* suborder (Figure 3A, B and Supplementary tables S10, S11). However, no signs of positive selection were found in the basal lineages of these suborders. As for the *Pantherine* family, only ZP2 exhibits diversifying selection effects (Figure 3A). Conversely, the subunits involved in cross-linking between ZP protein filaments ZP1 and ZP4 presented a notably lower prevalence of positive selection. No diversifying positive selection was detected in any of the analysed clades, and a positive selection signal was detected solely in the basal lineage of caniforms for the ZP4 protein (Figure 3C and Supplementary table S12). ZP1 did not display any selection signals in any of the analysed clades or branches (Figure 2D and Supplementary table S13).

**Figure 3:**
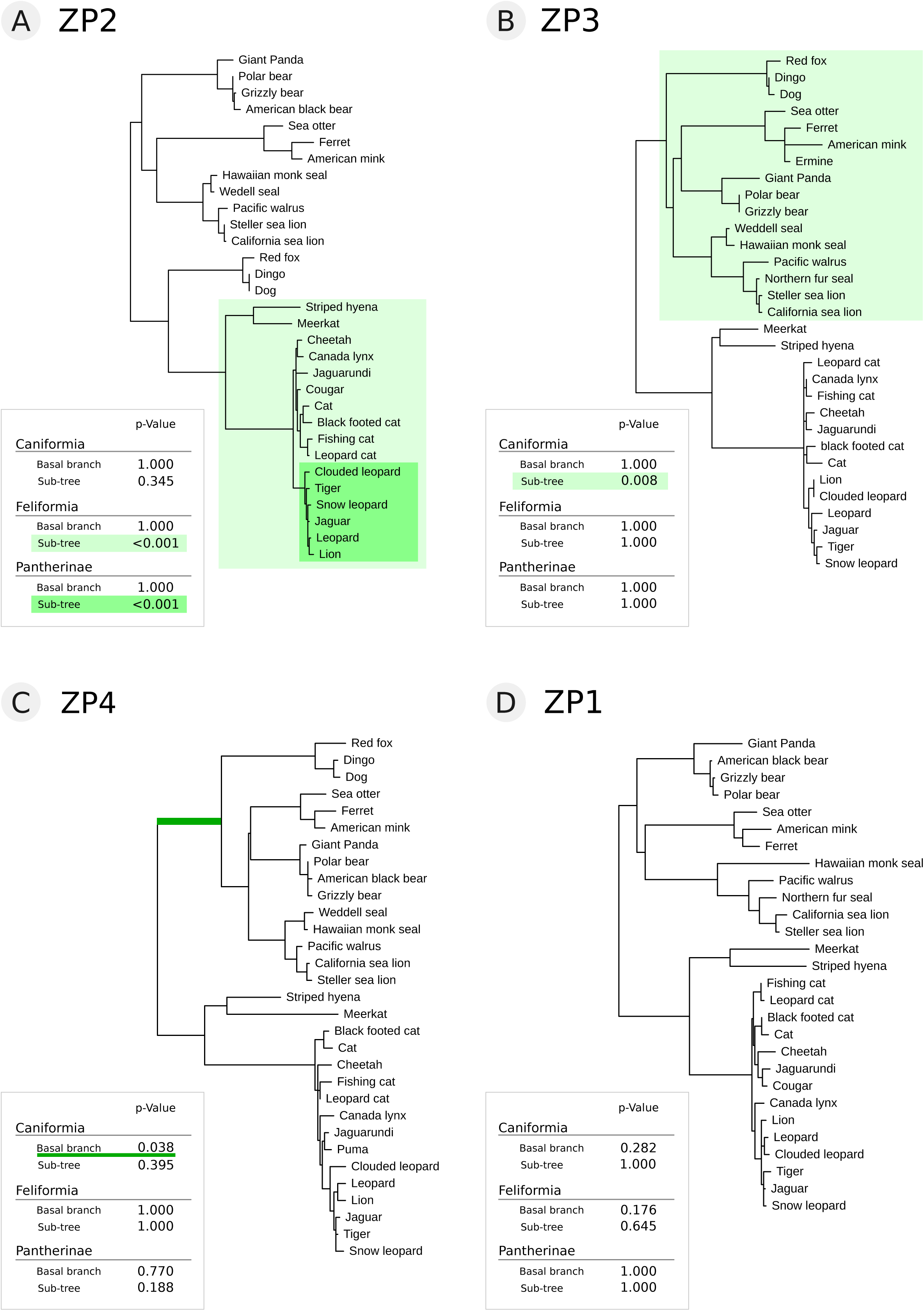
Phylogenetic trees and ZP proteins positive selection analyses results. Maximum likelihood phylogenetic reconstruction for ZP2 (A), ZP3 (B), ZP4 (C) and ZP1 (D) using 31 carnivore species in the cases of ZP2, ZP3 and ZP4 and 28 carnivore species for ZP1. Highlighted light green areas in A and B indicate divergent positive selection in *Feliformia* and *Caniformia* suborders in ZP2 and ZP3, respectively. The darker green highlighted area in A indicates divergent positive selection in the *Pantherinae* subfamily for ZP2. Dark green line in C indicates episodic positive selection in the *Caniformia* basal branch for ZP4. In the lower left corner of A, B, C and D panels tables summarize positive selection results for each protein in each of the basal lineages and subtrees tested.

### 3.4 ZP proteins Positive Selected Sites

ZP2 evolutionary analyses consistently revealed 2 candidate sites subjected to positive selection: jaguar positions 231 Q>V and 324 S>V/F/L, located in ZP-N2 and ZP-N3 respectively (Figure 4A, 4B and Supplementary figures S15, S16). Position 231 displays a glutamine (Q) for all caniform species and for some, but not all, feliform species. The remaining feliform species changed to a valine (V). While some feliform species conserve the glutamine, the feliform species that change to a valine involve different feliform families, including the pantherines. This indicates that this change occurred repeatedly and independently many times along the *Feliformia* suborder. On the other hand, position 324 is a serine (S) in all caniform species but changes to valine (V) and phenylalanine (F) in feliforms and to leucine (L) in some pantherine species (Figure 4C). It is important to mention that in jaguar ZP2 protein the sperm-binding region extends from position 55 to position 154 and ovastacine cleavage site appears between positions 171-172 (Figure 4A).

**Figure 4:**
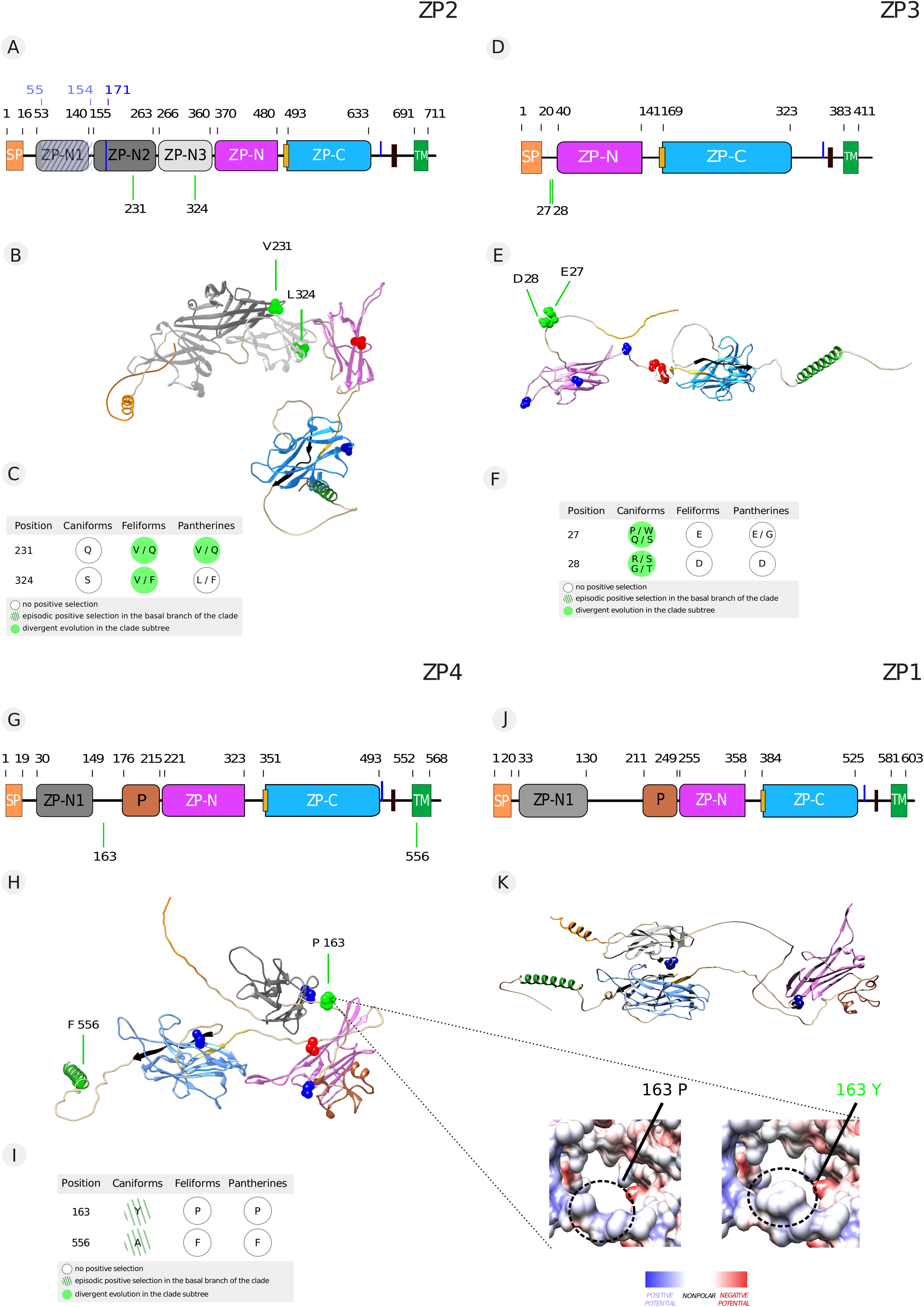
ZP proteins positive selected sites. A. Schematic representation of jaguar ZP2 protein domains. Idem Fig 1B. Light green lines below indicate positive selected sites. B. Three-dimensional model of the jaguar protein ZP2 shown in ribbons. N-glycosylated and O glycosylated sites are shown as blue and red atom spheres, respectively. Position and identities of positively selected amino acids are indicated and highlighted in light green atoms spheres. C. Table displaying sites under positive selection in ZP2 and their identities in the groups of interest. D. Schematic representation of jaguar ZP3 protein domains. Idem Fig. 1C. Light green lines below indicate positive selected sites. E. Three-dimensional model of the jaguar protein ZP3 shown in ribbons. N-glycosylated and O glycosylated sites are shown as blue and red atom spheres, respectively. Position and identities of positively selected amino acids are indicated and highlighted in light green atoms spheres. F. Table displaying sites under positive selection in ZP3 and their identities in the groups of interest. G. Schematic representation of jaguar ZP4 protein domains. Idem Fig. 1D. Light green lines below indicate positive selected sites. H. Three-dimensional model of the jaguar protein ZP4 shown in ribbons. N-glycosylated and O glycosylated sites are shown as blue and red atom spheres, respectively. Position and identities of positively selected amino acids are indicated and highlighted in light green atoms spheres. Coulombic surfaces depict the ancestral and derived states of positively selected amino acids, focusing solely on those sites that exhibit significant electrostatic charge changes from the ancestral to the derived state. I. Table displaying sites under positive selection in ZP4 and their identities in the groups of interest. J. Schematic representation of jaguar ZP1 protein domains. Idem Fig 1 E. K. Three-dimensional model of the jaguar protein ZP1 is shown in ribbons. N-glycosylated sites are shown as blue atom spheres.

The positive selection analysis of ZP3 revealed two sites that contribute to the diversifying selection signal within the *Caniformia* suborder. These sites are located at the N-terminal end of the protein and reside within a region that does not correspond to any known domain (Figure 4D and 4E). The adjacent sites are identified as follows: 27 E/G>S/W/Q/P and 28 D>R/T/G (Figure 4F and Supp. Figure S18, S19). Both sites exhibit acidic amino acids in feliforms, while in caniforms, they display a broader variability encompassing various amino acid types. Specifically, site 27 is a glutamic acid (E) found in most feliform species, except for the tiger and the snow leopard, which present a glycine (G). Conversely, among caniforms, site 27 exhibits a variety of amino acids, including serine (S), proline (P), and tryptophan (W). On the other hand, site 28 contains an aspartic acid (D) in all feliform species and displays variability among caniforms, where it can be either an arginine (R), glycine (G) or threonine (T) (Supplementary figures S18, S19). These sites are not located within any protein domain and, in the three-dimensional protein model, they are not found in proximity to N- or O-glycosylation sites (Figure 4E).

Moving on to the ZP subunits involved in cross-linking, sites under positive selection were only detected for ZP4. The single selection signal detected for this protein was particularly localized in the basal lineage of the caniforms (Figure 3C). Specifically, the changes detected in ZP4 for the basal caniform lineage were at positions 163 P>Y and 556 F>A (Figure 4I and Supplementary figure S21, S22). Site 163 is located in the N-terminal portion of the ZP4 protein, in an inter-domain region between the ZP-N1 domain and the trefoil domain (Figure 4G and 4H). In feliforms, this site is a proline (P), while in caniforms, it is a tyrosine (Y). Jaguar ZP4 protein structural model and disorder prediction data shows that site 163 is located in an unstructured and disordered proline rich region (Figure 4H and Supplementary figure S20). This site is spatially close to the N-glycosylated site at position 237, which belongs to the ZP-N domain of the ZP module (Figure 4H). Site 556 is a phenylalanine (F) in all feliforms and an alanine (A) in all caniforms. This site it is located at the beginning of the transmembrane region that anchors the immature protein to the oocyte membrane (Figure 4G, 4H). The entire C-terminal end portion of ZP proteins, including this transmembrane region, is cleaved to release the mature protein and allow its integration into the ZP. The C-terminal portion remains anchored to the membrane and is subsequently degraded. Moreover, both alanine and phenylalanine belong to the group of hydrophobic amino acids, characteristic of a transmembrane region. The 556 F>A change does not alter the nature of this amino acid, but it could produce a steric difference due to the size discrepancy between the lateral groups of these amino acids.

There are no positive selected sites to be reported in ZP1 the analyses showed an absence of positive selection in the basal lineages of *Caniformia*, *Feliformia*, and the *Pantherine* family, as well as an absence of diversifying positive selection within these clades (Figure 4J, 4K).

All domain arrangements for each subunit of jaguar ZP proteins were successfully predicted through AlphaFold modelling (Figure 4B, 4E, 4H, and 4K) (Supp figures S14, S17, S20, S23). These structural models were also used to spatially locate the N-glycosylated and O-glycosylated sites in an attempt to discover if they were positioned close to the detected selected sites as these sites are likely involved in ZP polymerization (Monné et al., 2008).

### 3.5 IZUMO1-JUNO positive selection analyses

Regarding the IZUMO1-JUNO interaction pair, an evolutionary study was carried out to find out which were the main evolutionary forces that shaped the history of each of the members of this pair involved in the fusion between gametes. The first to highlight is that, of all the proteins analysed in this work, IZUMO1 was the only one for which we were able to obtain predicted sequences with good quality for all the species of the *Carnivora* order. While JUNO also yielded a substantial number of coding sequences from *Carnivora* species, two sequences from Ermin (*Mustela erminea*) and the American black bear (*Ursus americanus*) were excluded due to sequencing quality issues. Despite these minor differences, assembling and curating both alignments proved relatively straightforward, with ample availability of coding sequences among *Carnivora* species and minimal annotation difficulties. However, as we reconstructed the phylogenetic trees for both genes, intriguing differences in their evolutionary histories began to emerge. The phylogenies of both genes yield the same general topology between clades of the *Carnivora* order. However, within the suborder Feliforma, it was not possible to make valid topology comparisons between trees beyond the split between species in the family *Felidae* and species belonging to other families of feliforms (Figure 5A). This limitation arose due to the low levels of branch support (<30%) obtained for JUNO along the whole *Felidae* family. While a few cases of low branch support were also detected in the *Felidae* family for IZUMO1, this phenomenon was significantly more noticeable in the case of JUNO, where it was impossible to draw confident conclusions about the topology of the *Felidae* family due to this limitation.

**Figure 5:**
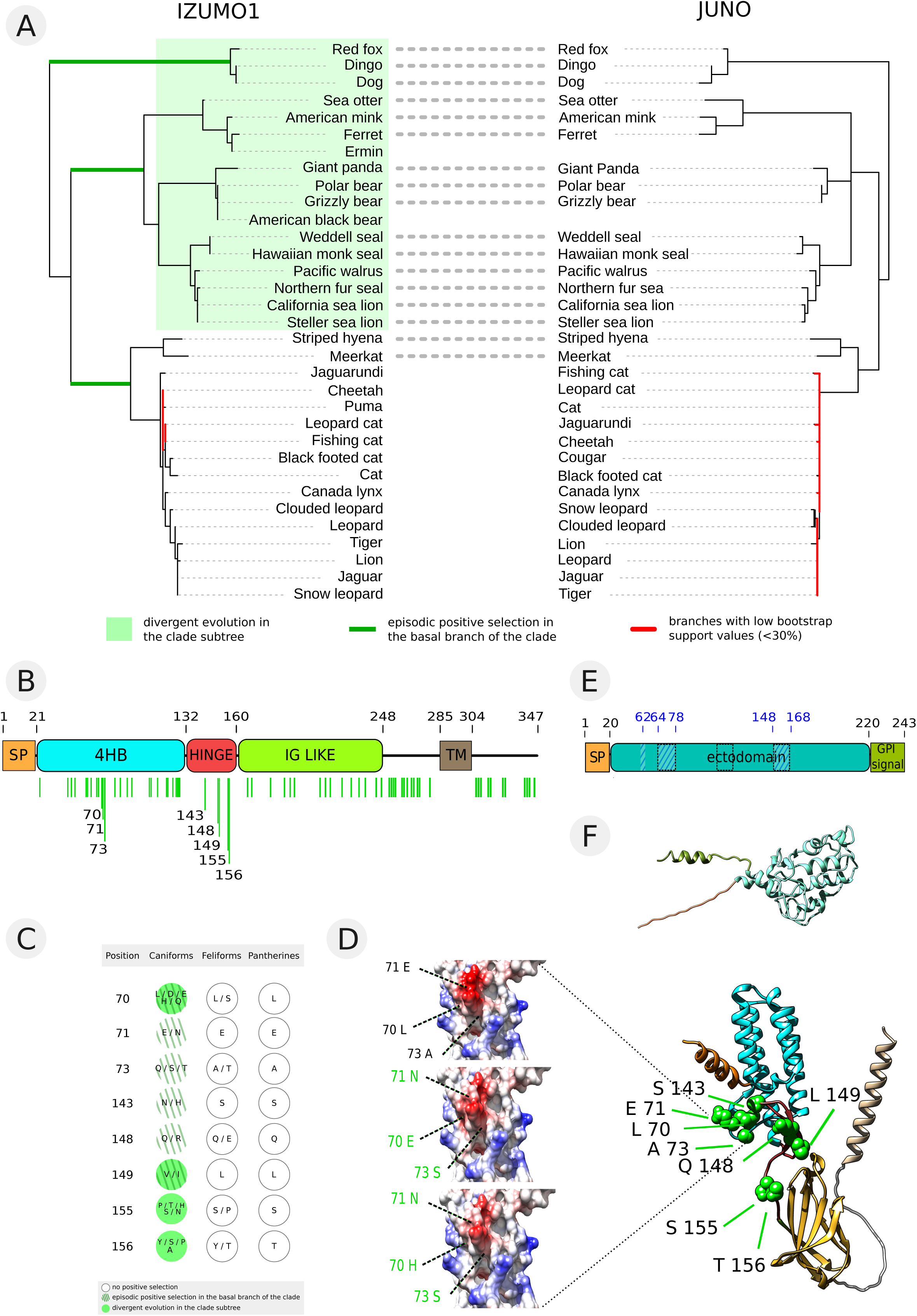
Evolution of IZUMO1-JUNO proteins fusion pair. A. Mirrored IZUMO1 and JUNO maximum likelihood trees showing comparable topologies (dotted lines). Episodic positive selection in the basal lineage of a group is indicated with dark green lines. Highlighted light green area indicates divergent positive selection. Branches with bootstrap values below 30% are highlighted in red. B. Schematic representation of jaguar IZUMO1 protein domains. Idem Fig. 1F. Light green lines below indicate positive selected sites. C. Table displaying sites under positive selection in IZUMO1 and their identities in the groups of interest. Only positive selected sites previously reported to be involved in interaction with JUNO are shown. D. Three-dimensional model of IZUMO1 protein shown in ribbons. Position and identities of positively selected amino acids are indicated and highlighted in light green atoms spheres. Coulombic surfaces depict the ancestral and derived states of positively selected amino acids, focusing solely on those sites that exhibit significant electrostatic charge changes from the ancestral to the derived state. E. Schematic representation of jaguar JUNO protein domains. Idem Fig. 1G. F. Three-dimensional model of the jaguar protein JUNO shown in ribbons.

Significant differences emerged from the positive selection analyses of IZUMO1 and JUNO. The evolutionary analysis of IZUMO1 revealed signatures of adaptive evolution in the basal lineages of both *Caniformia* (p-value = 5.12e^-10^) and *Feliformia* (p-value = 0.0366) suborders, but not in the basal *Pantherinae* lineage. Moreover, strong diversifying positive selection marks were detected within the *Caniformia* suborder (p-value = 2.18e^-17^) (Table 1). It is worth noting that selection signatures found in IZUMO1, both episodic and diversifying, were significantly more intense in the *Caniformia* suborder than in the *Feliformia* basal lineage (Table 1). It is also important to highlight that such selection signatures were absent in the *Feliformia* suborder or the *Pantherine* subfamily (Figure 5A).

**Table 1.**
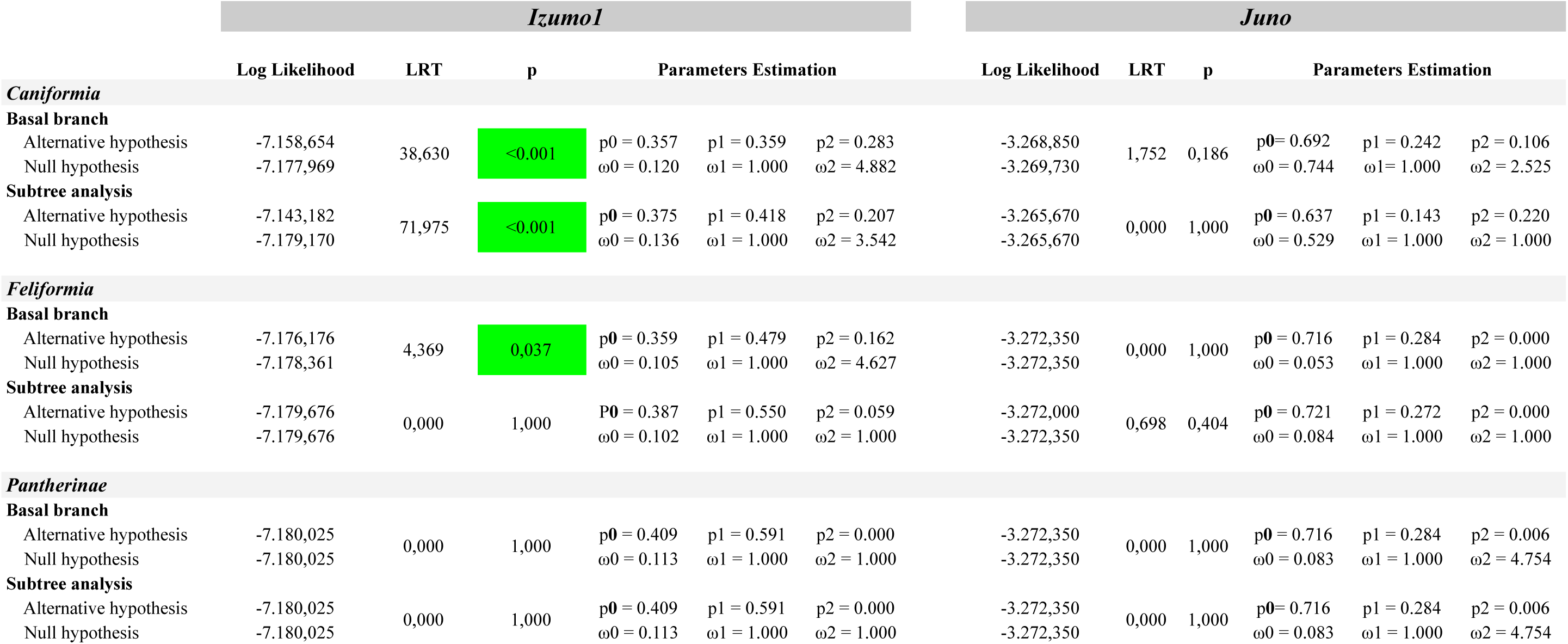
Branch-site positive selection results for *Izumo1* and *Juno* in the *Carnivora* order. Note: LRT: statistical value of the likelihood ratio test; p_0_-p_2_ and ω_0_-ω_2_ proportion of sites and dN/dS values estimated for each site class in the selection model (alternative hypothesis), respectively. Green highlighted p values show positive test results (p<0.05)

In contrast to the abundance of positive selection results for IZUMO1, the evolutionary analysis of JUNO revealed a complete absence of selection signals across all clades and basal lineages analysed (Figure 5A, 5E, 5F, Table 1 and Supplementary figure S28).

The results of the positive selection analyses for IZUMO1 revealed a considerable number of positively selected sites, precisely 69 sites (Figure 5B and Supplementary table S26). The majority of these sites correspond to adaptations in the basal lineage of caniforms and sites that are undergoing diversifying evolution within the *Caniformia* suborder. Only three of these 69 sites were detected in the basal lineage of feliforms. These sites are: 87 S/A>V; 181 R>K; 268 L/H/Q>T (Supplementary figure S25). This difference in the number of detected sites between the *Caniformia* and *Feliformia* suborders is not surprising when we contrast the resulting selection intensity values for these clades. The p-values when testing the hypothesis of selection against a null model for the basal lineage (indicative of adaptation through episodic selection) and the subclade (indicative of diversifying selection within the clade) of the *Caniformia* suborder were 5.12e^-10^ and 2.18e^-17^, respectively. In contrast, the p-value for the hypothesis test that detected positive selection in the basal lineage of the *Feliformia* suborder was 0.0366 (Table 1).

### 3.6 IZUMO1 Positive Selected Sites

Positively selected sites in IZUMO1 are uniformly distributed throughout the protein, with two important exceptions—the signal peptide and the transmembrane domain—which appear devoid of positively selected sites (Figure 5B). Several of the many IZUMO1 sites that were identified as having changed under the influence of positive selection in *Caniformia* suborder correspond to sites reported as important for the interaction with JUNO (Kato et al., 2016). Given the large number of detected sites for IZUMO1, we will focus only on those (the complete list of IZUMO1 positive selected sites can be found in Supplementary table S26). These sites can be divided into two clusters of closely located sites (Figure 5B and Supplementary figure S24). The first cluster includes sites 70, 71, and 72, situated in the 4HB domain, the four alpha helices bundle of the N-terminal region of the protein. The second cluster is located in the HINGE domain and comprises sites 143, 148, 149, 155, and 156.

Site 70 is a leucine (L) in all feliforms, except for the meerkat (*Suricata suricata*), which displays a serine (S). In contrast, caniforms show much more variability at this site, changing to aspartic acid (D) in bears, glutamic acid (E) in pinnipeds, and histidine (H) in mustelids (Supplementary figure S25). These changes involve the transition from a hydrophobic amino acid to variants with charge (Supplementary figure S27).

Adjacent to the previous site is site 71 E>N. This site contains a glutamic acid (E) in almost all species of the *Carnivora* alignment, except for species in the Canidae family (Supplementary figure S25). These species exhibit an asparagine (N), leading to a change from a negatively charged amino acid to an uncharged polar one (Supplementary figure S27). The electrostatic charge variation induced by the two aforementioned changes can be visualized on the Coulombic surface in figure 5D. Site 73 A>T/S/Q presents an alanine (A) in all species of the *Felidae* family and a threonine (T) in the striped hyena (*Hyaena hyaena*) and meerkat (*Suricata suricata*), feliform species outside the *Felidae* family. On the other hand, in caniforms it shows a glutamine (Q) for species in the Canidae family and a serine (S) in all other caniform species (Supplementary figure S25).

Regarding the second cluster of positive selected sites crucial for the interaction of IZUMO1 with its counterpart JUNO, we can start with site 143 S>N/H. This site shows a serine (S) in all feliform species and an asparagine (N) in the majority of caniform species (Supplementary figure S25). This change may not be highly significant as both amino acids have uncharged polar side chains. However, in species of the Canidae family, this position shows a histidine (H), a positively charged amino acid (Supplementary figure S27).

Site 148 Q>R is a glutamine (Q) for most of the species in the alignment, both caniforms and feliforms. While it remains almost constant in feliforms, changing only in the meerkat (*Suricata suricata*), in caniforms, it changes to an arginine (R) in the ferret (*Mustela putorius*) and in canid species (Supplementary figure S25).

Site 149 L>V/I is a leucine (L) in all feliforms, but it changes to a valine (V) in caniforms and then again to an isoleucine (I) in canid species (Supplementary figure S25).

Sites 155 and 156 are adjacent and both are under the effects of diversifying positive selection. Site 155 S>P/T/H/S is a serine (S) in almost all feliform species, except for the meerkat, while in caniforms, it shows significant variability between serine (S), histidine (H), proline (P), and threonine (T) (Supplementary figure S25). Lastly, we have site 156 T/Y>A/P/S/Y. This site shows a threonine (T) in all felid species and a tyrosine (Y) in all non-felid feliform species. In caniforms, this site also displays substantial variability between alanine (A), proline (P), serine (S), and tyrosine (Y) (Supplementary figure S25).

A general summary of the positive selection results for all the proteins studied across all analysed clades can be observed in figure 6.

**Figure 6:**
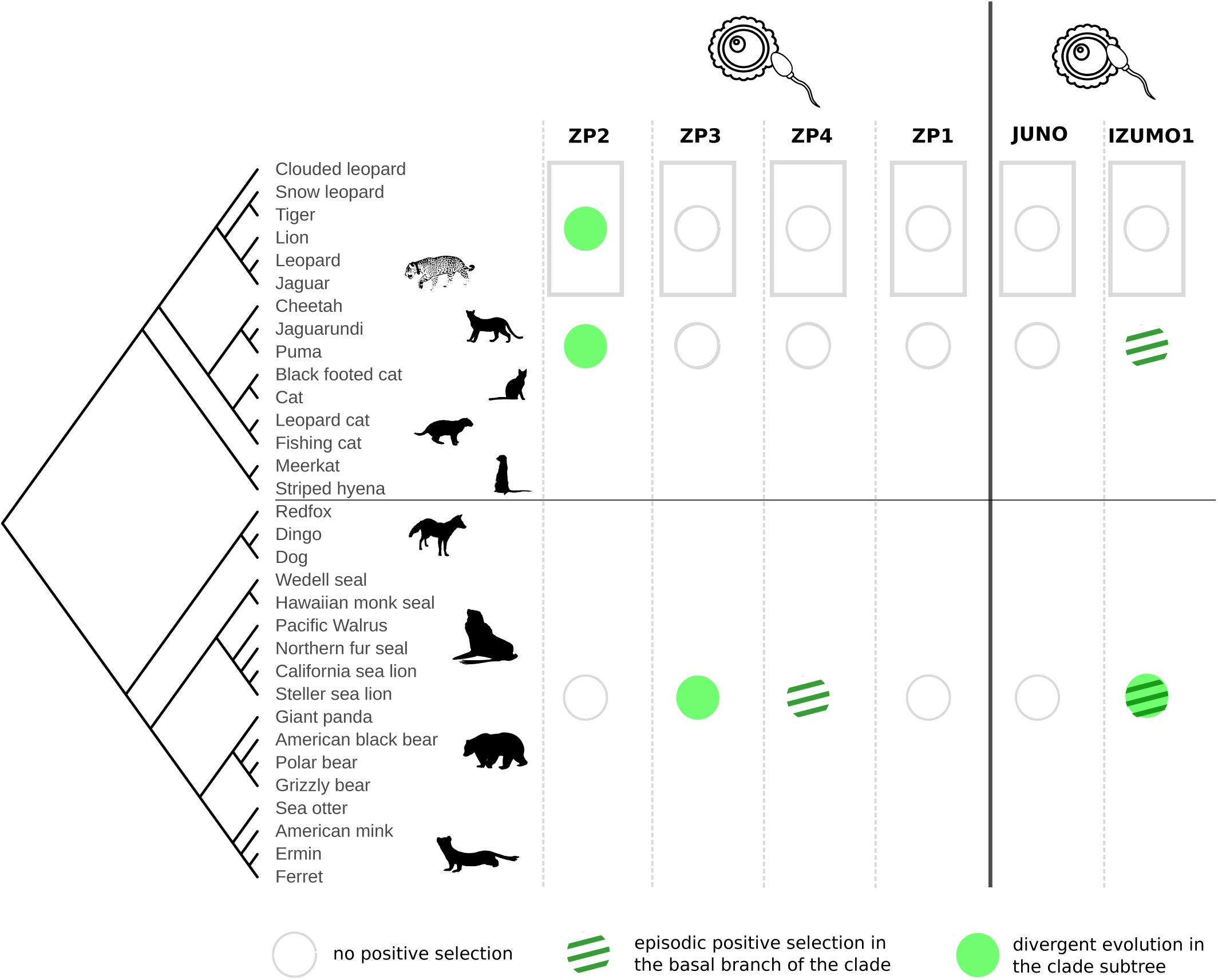
Summary of Positive selection analyses. Positive selection analyses results for ZP proteins, JUNO and IZUMO1 in all tested groups: caniforms, feliforms and pantherines. A filled circle with green diagonal stripe pattern indicates episodic positive selection at the basal branch of the clade. A light green circle indicates divergent positive selection in the clade sub-tree. Empty white circle denotes absence of positive selection. Species tree is representative of the full set of carnivore species employed in this study.

### 3.7 Control proteins: ACTN1, a neutrally evolving protein, and TPM3, a jaguar positively selected protein

To demonstrate that the likelihood methods do not spuriously detect selection, we performed likelihood ratio tests for the conserved housekeeping *Actn1* gene (Holterhoff et al., 2019) (a negative control protein). Supplementary table S29 shows the negative expected results.

To demonstrate the reliability of the method used to detect positive selection and inferring sites potentially subjected to selection, we performed likelihood ratio tests on control proteins, using *Tpm3* one of the positively jaguar specific selected genes previously reported (Figueiró el al., 2017). Results shown in Supplementary table S30 shows a p-value=0.0006.

## 4. Discussion

This study sheds light the evolution of ZP proteins and the IZUMO1-JUNO gamete interaction pair within the order *Carnivora*. The rapid evolution of reproductive proteins can make a significant contribution to reproductive isolation between diverging taxa, given the possibility that changes in these proteins may lead to isolation between species (Swanson et al., 2001; Swanson & Vacquier, 2002; Turner & Hoekstra, 2008). In order to assess this in the *Carnivora* order, a comprehensive compilation of all available coding sequences from orthologs across species within this order was required. As a result, the most complete and curated collection of aligned carnivorans sequences for these fertilization-related proteins to date was obtained. To the best of our knowledge, no study has so far conducted such a meticulous and detailed investigation into the evolution of these gamete-interaction proteins in the *Carnivora* order. Initially, sequence identity analyses were carried out to characterize and compare the amino acid diversity between these gamete-interaction proteins, as well as to evaluate differences between clades within the *Caniformia* suborder. Subsequently, site-lineage-specific analyses of positive selection were performed to detect signals of molecular adaptation and diversifying evolution. These selection hypotheses were tested for both *Carnivora* suborders, *Caniformia* and *Feliformia*, as well as the *Pantherinae* subfamily. In cases where sites under positive selection were identified, these were mapped onto structural models of these proteins generated for jaguar sequences. This allowed us to gain insights into the evolutionary forces acting upon gamete interaction proteins in the different clades of the *Carnivora* order. The results obtained in this study revealed that the *Feliformia* suborder exhibits significantly lower diversity than its sister taxa, *Caniformia*, across all examined proteins. Regarding ZP2 and ZP3 molecular evolution, essential ZP protein family subunits directly involved in sperm recognition, the presence of diversifying positive selection in the *Feliformia*, *Caniformia*, and *Pantherinae* clades was identified. However, no relevant changes could be linked to sperm binding. Concerning the IZUMO1-JUNO fusion pair, IZUMO1 exhibited the highest number of positive selection signals and sites in both *Caniformia* and *Feliformia* basal lineages and also along the *Caniformia* subtree, but not in the *Pantherinae* basal lineage or subtree. On the other hand, JUNO showed an absence of any signatures of positive selection across all tested lineages and clades. Within the positively selected sites of IZUMO1, eight were identified, clustering in two regions involved in the interaction with JUNO.

Multiple driving forces have been proposed to explain the occurrence of positive selection in reproductive loci involved in speciation. One prominent hypothesis is sperm competition, which refers to the competition between sperm from different males for the fertilization of a female’s eggs. This theory posits that sperm competition instigates a continuous and adaptive “arms race,” wherein the selective intensity is expected to correlate with the degree of polyandry (Clark et al., 2006). Under this hypothesis, male proteins involved in various aspects of fertilization, including locating, reaching, binding, penetrating, and fusing with the egg, may undergo adaptive evolution.

On the other side, given the larger energy investment in female gametes, polyspermy can have more detrimental effects on female fitness than male fitness. Consequently, female gamete proteins are expected to evolve to lower the fertilization rate, while sperm proteins continuously strive to increase it within a competitive context. This leads to the concept of sexual conflict over adaptive optima, whereby females and males engage in counter-adaptations, resulting in a co-evolutionary chase between male and female characters (Swanson & Vacquier, 2002). This conflict may maintain either polymorphism or extend a co-evolutionary competition between egg and sperm proteins (Turner & Hoekstra, 2008). Sperm competition, which promotes rapid rates of fertilization, may be advantageous for males, whereas females may benefit from a more moderate fertilization rate to prevent polyspermic fertilization. In addition to sexual conflict and sperm competition, sexual selection plays a crucial role in shaping mating behaviour and displays. It is conceivable that sexual selection also operates at the gametic level, wherein an egg exhibits a preference for a specific sperm protein allele, resulting in assortative mating (Eberhard, 1996). Another driving force is reinforcement, which refers to the evolution of reproductive barriers to prevent hybridization. When allopatric populations hybridize and produce offspring with reduced fitness, reinforcement explains the divergence observed in gamete recognition proteins (Swanson & Vacquier, 2002). While the aforementioned forces primarily stem from endogenous factors within the reproductive system of the species, it is important to consider exogenous forces as well. Pathogen resistance, for instance, represents an exogenous force that may drive divergence at these reproductive loci. Microbial attacks exert constant pressure on gamete surface proteins, necessitating their continual adaptation to evade these attackers (Vacquier et al., 1997). It is important to highlight that several of these hypotheses share overlapping predictions, rendering their differentiation challenging (Clark et al., 2006; Carlisle & Swanson, 2021).

### Characterization of jaguar gamete interaction proteins

All jaguar sequences for ZP proteins and the IZUMO1-JUNO fusion pair obtained during this study show a high identity levels with sequences from other species of the *Pantherinae* subfamily. This effect was also observed to some extent in the suborder *Feliformia*, but not in the sister suborder *Caniformia*. In terms of locus structure preservation, each of the studied genes presents a high consistency in number of coding exons and their distribution and grouping along the locus for all feliform species. This aligns with the high levels of genomic stability reported for the felid lineage (Bredemeyer et al., 2023). The only exception is the species of the puma taxonomic group (*P. concolor* and *H. yagouaroundi*), that present a higher locus structural variability.

### ZP proteins evolution

*A priori*, it might be proposed that reproductive genes under positive selection would be expected to play secondary roles in reproductive mechanisms, while proteins essential for fertilization would generally remain conserved. However, positive selection has been detected numerous times in reproductive proteins, challenging this expectation (Grayson et al., 2015). The findings presented in this study support this apparent contradiction. Essential ZP proteins subunits directly involved in fertilization and oocyte interaction, ZP2 and ZP3, display positive selection signatures that do not involve changes in sperm recognition regions demonstrated in other model organisms. On the other hand, the subunits that play a secondary role structuring the ZP matrix, ZP1 and ZP4, exhibit lower or even no levels of positive selection. Moreover, ZP1 and ZP4 are the two subunits of the ZP protein family that appear pseudogenized in some mammalian species (Moros-Nicolás et al., 2018a; Goudet et al., 2008; Lefièvre et al., 2004) consistent with the fact that fertilization can still occur even in their absence, possibly due to a functional redundancy between them.

ZP2 positive selected sites, 231 Q>V and 324 S>V/F, in the jaguar would not affect ZP2 sperm-recognition region reported as essential in mice (Gahlay et al., 2010; Avella et al., 2014; 2016). After fertilization the ZP2 N-terminal region is cleaved by the ovastacine metalloproteases in order to prevent polyspermy (Burkart et al., 2012). Positive selected amino acids in ZP2 do not affect ovastacine cleavage site in feliforms or pantherines, as this sequence appears in the jaguar sequence between positions 171-172. However, as they are located in N-terminal ZP-N domains, in ZP-N2 and ZP-N3, respectively, they may be involved in the interaction between ZP filaments (Jovine et al., 2002; 2005).

Both positive selected sites revealed for ZP3, 27 E/G>S/W/Q/P and 28 D>R/T/G, are adjacently located at the N-terminal end of the mature protein. These sites hold the strongest diversifying selection signal for caniforms in this protein. In feliforms, both sites show acidic amino acids, while in caniforms, they exhibit a wider range of variability, including various amino acid types. This change in charge and the N-terminal location of these amino acids could indicate that they vary in the suborder *Caniformia* due to an external factor of selective pressure, such as sperm competition, sexual conflict, sexual selection, or escape from pathogens. However, the location of these sites in the N-terminal extreme of ZP3 protein and outside any domain or functional region would indicate that they are not involved in ZP3 function.

The selection signature detected for ZP4 is specifically located in the basal lineage of the *Caniformia* suborder. This, unlike the selection signatures detected for ZP2 and ZP3, suggests an episodic process that promoted the appearance of variants that were subsequently conserved. This means that detected adaptive changes (163 P>Y and 556 F>A) in this lineage occurred early in the history of this suborder, after their split from *Feliformia*, and were fixed by selection. Taking into account that these changes took place in the early history of the *Caniformia* suborder and were later fixed, they may be involved in the surge of a prezygotic isolating mechanism between caniforms and its sister group *Feliformia*. The change 163 P>Y is more relevant than 556 F>A, which is not present in the mature protein and keeps the same amino acid hydrophobic nature. Site 163 is located in the unstructured and disordered region between ZP-N1 and trefoil domains. The accumulation of prolines in this inter-domain sequence provides stiffness to this linker region (Krieger et al., 2005). The change 163 P>Y would reduce this stiffness and increases the mobility of the region and between the adjacent domains.

ZP1 was the only ZP protein that showed no signatures of positive selection in any of the lineages or clades tested. Although ZP1 and ZP4 remain by far the less studied components of the mammalian ZP, Jovine’s lab structurally characterized ZP1 cross-linking function in human proteins (Monné & Jovine, 2011; Nishimura et al., 2019). Studies of human ZP1 mutant protein suggest that lack of a ZP in infertile patients carrying the corresponding mutation could be due to impaired secretion of ZP1 and the consequent absence of ZP filament cross-linking. As this step is prior to the cross linking with ZP3 it cannot discriminate how this region of ZP1 would interact with ZP3 if at all (Huang et al., 2014). On the other hand, the trefoil/ZP module region of ZP1 does not form non-covalent homodimers in humans, which shows that ZP1-N1 necessarily participates in cross-linking to maintain human ZP with its consequences over fertility (Nishimura et al., 2019).

### IZUMO1-JUNO evolution

Sequence identity analyses revealed that the sperm protein IZUMO1 displays significantly higher variability than the oocyte proteins. It is likely that this difference between sperm proteins and oocyte proteins is due to sperm competition. This would explain the widespread diversifying selection detected for IZUMO1. On the other side, its oocyte partner, JUNO, rendered no signature of positive selection in any of the lineages or clades tested. However, it displays variability comparable to other oocyte proteins. Therefore, the absence of positive selection signals in JUNO is not due to an extremely conserved nature of this protein but rather to a strictly neutral evolution process. Similar strikingly differences in evolutionary forces had been previously identified acting on gamete interaction proteins in abalone Lysin-VERL pair (Swanson et al., 2001). However, for the mammalian IZUMO1-JUNO pair a coordinated evolution hypothesis has been proposed given that both proteins experience positive selection in mammals, although these signals appear in different clades for each partner (Grayson et al., 2015). The phylogenetic scope used in the present work is tighter, as it focuses on the *Carnivora* order, and results obtained for JUNO indicate a total absence of positive selection in this group.

A recent article reported that the tryptophan at position 148 of IZUMO1 was crucial for the binding with JUNO (Brukman et al., 2023). This position is not affected by positive selection and is completely conserved in all *Carnivora* species. All 8 IZUMO1 positive selected sites detailed here are also known to be involved in the interaction with JUNO (Kato et al., 2016). Therefore, they may be used to generate mutations in future functional studies seeking to gain insight in IZUMO1-JUNO binding in carnivorans.

### Final conclusions

Here we reported 8 IZUMO1 positive selected sites, all of them obtained from the selection analyses of the *Caniformia* basal lineage and subtree, which are also involved in JUNO interaction. No other positively selected site that appears to be directly involved in sperm-egg binding was found in the remaining gamete-interacting proteins. ZP2 changes observed for feliforms and pantherines may be affecting the interaction of this protein with other filaments that structure the ZP matrix, but they do not reside in the known region for sperm interaction as described in other mammalian models. ZP3 positively selected sites detected in caniforms are not located within any functional domains of this protein. Lastly, ZP4 sole detected site that is present in mature protein seems to influence the flexibility of an interdomain region. ZP1 and JUNO, on the other hand, exhibited a complete absence of positively selected sites in carnivores.

The main limitation of this work lay in the difficulty of establishing a functional role for the detected positive selected sites. Another significant limitation is reliably discerning the origin of positive selection. While this process is typically associated with molecular adaptation in a traditional way, there are various hypotheses explaining its occurrence in reproductive proteins, and distinguishing between them poses a challenging task. Although it is still widely believed that positive selection directly influences egg-sperm recognition, we do not exclude the possibility that positive selection may have in fact an indirect role. Experimental evidence comes from VERL/lysin complexes in which positive selection at the level of VERL repeat 1 (and to a lesser extent VERL repeat 2) generates a binding affinity gradient for lysin that results in species-specificity (Raj et al., 2017). However, this is not done by directly affecting residues at the interface. In fact, the divergence of repeat 1 may even be due to completely different reasons - for example pathogen avoidance (Raj et al., 2017).

In total, there is evidence suggesting a lack of strong interspecific isolation barriers in pantherines (Li et al., 2016; Figueiró et al., 2017; Yadav et al., 2019; Duque Rodriguez et al., 2021). The absence of positive selection for this lineage in gamete recognition regions of reproductive proteins reported here is consistent with the premise of ancient and ongoing hybridization between some pairs of big cat species. Given that successful fertilization relies on the molecular compatibility between gametes, as long as sperm and egg can recognize and bind via their surface proteins, gamete fusion is possible between individuals of different species. This would be permissive of hybridization events possibly impacting speciation processes (Gert et al., 2023). From a conservation perspective it also means that declining small populations like jaguars in Argentina could be susceptible to hybridization, though on the other hand future efforts to enrich local gene pools by assisted migration from different conspecific populations - if considered necessary - would not be hampered by gamete incompatibility (Hoffmann et al. 2021; Johnson et al. 2010).

## Author Contributions

CC, CP and FP performed the bioinformatic analyses. FP designed the evolution study. CDB, FP, CC and CP performed jaguar ZP proteins structural models. TG contributes with phylogenic approaches and all concerning Vagalume jaguar genome data. PS conceived the project. FP and PS wrote the paper. All authors discussed the results and commented on the manuscript.

## Acknowledgments

We thank DNAZoo, particularly Olga Dudchenko and Erez Lieberman-Aiden for discussion about HiC jaguar genome data, Luca Jovine for fruitful paper and idea discussions. We also thank Fundación Williams, FIByME for the administration of grants and aided the author’s work.

## Data Availability

The data that support the findings of this study are openly available in figshare at http://doi.org/10.6084/m9.figshare.23896791.v3, reference number 23896791.

This supporting data includes: gene sequence alignments, phylogenetic trees and positive selection analyses files for all tested proteins in this work.

