## Supplementary for "Positive selection in gamete interaction proteins in *Carnivora*"

MOLECULAR ECOLOGY

**TABLE OF CONTENTS:**


| | | |
|---|---|---|
| **Supplementary table S1** | origin of Panthera onca nucleotidic sequence | page 2 |
| **Supplementary table S2** | Zp2 accession numbers | page 3 |
| **Supplementary table S3** | Zp3 accession numbers | page 4 |
| **Supplementary table S4** | Zp4 accession numbers | page 5 |
| **Supplementary table S5** | Zp1 accession numbers | page 6 |
| **Supplementary table S6** | Izumo1r accession numbers | page 7 |
| **Supplementary table S7** | Izumo1 accession numbers | page 8 |
| **Supplementary table S8** | pairwise sequence identities report for Carnivora in each studied protein | page 9 |
| **Supplementary table S9** | pairwise sequence identities report for taxonimic groups of interest (feliformia, caniformia and pantherinae) in each studied protein | page 10 |
| **Supplementary table S10** | branch-site positive selection results for Zp2 | page 11 |
| **Supplementary table S11** | branch-site positive selection results for Zp3 | page 12 |
| **Supplementary table S12** | branch-site positive selection results for Zp4 | page 13 |
| **Supplementary table S13** | branch-site positive selection results for Zp1 | page 14 |
| **Suplementary figure S14** | secondary structure, topology, and DISOPRED analysis plot of the ZP2 protein | page 15 |
| **Suplementary figure S15** | detail of alignment with positively selected sites in ZP2 | page 16 |
| **Suplementary figure S16** | detail of coulumbic surface in ZP2 | page 17 |
| **Suplementary figure S17** | secondary structure, topology, and DISOPRED analysis plot of the ZP3 protein | page 18 |
| **Suplementary figure S18** | detail of alignment with positively selected sites in ZP3 | page 19 |
| **Suplementary figure S19** | detail of coulumbic surface in ZP3 | page 20 |
| **Suplementary figure S20** | secondary structure, topology, and DISOPRED analysis plot of the ZP4 protein | page 21 |
| **Suplementary figure S21** | detail of alignment with positively selected sites in ZP4 | page 22 |
| **Suplementary figure S22** | detail of coulumbic surface in ZP4 | page 23 |
| **Suplementary figure S23** | secondary structure, topology, and DISOPRED analysis plot of the ZP1 protein | page 24 |
| **Suplementary figure S24** | secondary structure, topology, and DISOPRED analysis plot of the IZUMO1 protein | page 25 |
| **Suplementary figure S25** | (a, b, c, d , e) detail of alignment with positively selected sites in IZUMO1 | pages 26-31 |
| **Supplementary table S26** | sites under positive selection in IZUMO1 | page 32 |
| **Supplementary figure S27** | detail of coulumbic surface in IZUMO1 | page 33 |
| **Suplementary figure S28** | secondary structure, topology, and DISOPRED analysis plot of the JUNO protein | page 34 |
| **Supplementary table S29** | branch-site positive selection results for control proteins using Actn1 | page 35 |
| **Supplementary table S30** | branch-site positive selection results for control proteins using Tpm3 | page 36 |

| **Supplementary table S1: origin of *Panthera onca* nucleotidic sequence** | | | |
|---|---|---|---|
| **gen** | **query sequence** | **database** | **length of the retrieved sequence (nucleotides)** |
| *Zp2* | *Panthera pardus* *XM_019430284.1* | *Panthera onca* chromosome-resolution Hi-C genome assembly 2 | 2151 |
| *Zp3* | *Panthera pardus* *XM_019421632.1* | *Panthera onca* chromosome-resolution Hi-C genome assembly 2 | 1272 |
| *Zp1* | *Panthera pardus* *XM_019414554.1* | Illumina *Panthera onca* genome assembly 1 | 1821 |
| *Zp4* | *Panthera pardus* *XM_019419088.1* | Illumina *Panthera onca* genome assembly 1 | 1704 |
| *Izumo1r* | *Panthera pardus* *XM_019450101.1* | *Panthera onca* chromosome-resolution Hi-C genome assembly 2 and Illumina *Panthera onca* genome assembly 1 | 729 |
| *Izumo1* | *Panthera leo* *ENSPLOG00000009340* | *Panthera onca* chromosome-resolution Hi-C genome assembly 2 | 1037 |

1: Figueiró et al., 2017
2: DNAZoo

**Supplementary table S2: *Zp2* accession numbers**

| | Species | | Accession number |
|---|---|---|---|
| **1** | Red fox | *Vulpes vulpes* | XM_026003244.1 |
| **2** | Dog | *Canis lupus familiaris* | NM_001003304.2 |
| **3** | Dingo | *Canis lupus dingo* | XM_035717942.1 |
| **4** | Grizzly bear | *Ursus arctos horribilis* | XM_044390073.1 |
| **5** | American black bear | *Ursus americanus* | XM_045776421.1 |
| **6** | Polar bear | *Ursus maritimus* | XM_008697192.2 |
| **7** | Panda | *Ailuropoda melanoleuca* | XM_002927601.4 |
| **8** | Steller sea lion | *Eumetopias jubatus* | XM_028095252.1 |
| **9** | Hawaiian monk seal | *Neomonachus schauinslandi* | XM_021692216.1 |
| **10** | California sea lion | *Zalophus californianus* | XM_027612146.1 |
| **11** | Pacific walrus | *Odobenus rosmarus* | XM_004415752.1 |
| **12** | Weddell seal | *Leptonychotes weddellii* | XM_006738322.2 |
| **13** | Sea otter | *Enhydra lutris* | XM_022501447.1 |
| **14** | Ferret | *Mustela putorius furo* | XM_004780136.2 |
| **15** | American mink | *Neovison vison* | XM_044233004.1 |
| **16** | Cat | *Felis catus* | NM_001009875.1 |
| **17** | Leopard | *Panthera pardus* | XM_019430284.1 |
| **18** | Tiger | *Panthera tigris* | XM_007084109.2 |
| **19** | Cheetah | *Acinonyx jubatus* | XM_027042947.1 |
| **20** | Jaguar | *Panthera onca* | - |
| **21** | Lion | *Panthera leo* | - |
| **22** | Snow leopard | *Panthera uncia* | - |
| **23** | Clouded Leopard | *Neofelis nebulosa* | - |
| **24** | Jaguarundi | *Herpailurus yagouaroundi* | - |
| **25** | Black footed cat | *Felis nigripes* | - |
| **26** | Fishing cat | *Prionailurus viverrinus* | - |
| **27** | Leopard cat | *Prionailurus bengalensis* | - |
| **28** | Lynx | *Lynx canadensis* | XM_030300094.1 |
| **29** | Puma | *Puma concolor* | XM_025916779.1 |
| **30** | Striped hyena | *Hyaena hyaena* | XM_039234420.1 |
| **31** | Meerkat | *Suricata suricatta* | XM_029947959.1 |

**-** : nucleotide sequences obtained through customized local implementation of BLAST.

**Supplementary table S3: *Zp3* accession numbers**

| | Species | | Accession number |
|---|---|---|---|
| **1** | Hawaiian monk seal | *Neomonachus schauinslandi* | XM_021700471.1 |
| **2** | Northern fur seal | *Callorhinus ursinus* | XM_025864635.1 |
| **3** | California sea lion | *Zalophus californianus* | XM_027612093.2 |
| **4** | Steller sea lion | *Eumetopias jubatus* | XM_028110131.1 |
| **5** | Weddell seal | *Leptonychotes weddellii* | XM_006733910.1 |
| **6** | Pacific walrus | *Odobenus rosmarus* | XM_004399076.2 |
| **7** | American mink | *Neovison vison* | XM_044233042.1 |
| **8** | Sea otter | *Enhydra lutris* | XM_022501525.1 |
| **9** | Ferret | *Mustela putorius furo* | NM_001310185.1 |
| **10** | Ermine | *Mustela erminea* | XM_032326901.1 |
| **11** | Polar bear | *Ursus maritimus* | XM_008698535.2 |
| **12** | Panda | *Ailuropoda melanoleuca* | XM_002925817.4 |
| **13** | Grizzly bear | *Ursus arctos horribilis* | XM_026516370.2 |
| **14** | Dingo | *Canis lupus dingo* | XM_025426327.2 |
| **15** | Dog | *Canis lupus familiaris* | NM_001003224.1 |
| **16** | Red fox | *Vulpes vulpes* | XM_026019513.1 |
| **17** | Tiger | *Panthera tigris* | XM_007073897.2 |
| **18** | Leopard | *Panthera pardus* | XM_019421632.1 |
| **19** | Cat | *Felis catus* | NM_001009330.2 |
| **20** | Jaguar | *Panthera onca* | - |
| **21** | Cheetah | *Acinonyx jubatus* | XM_015078784.2 |
| **22** | Lion | *Panthera leo* | - |
| **23** | Snow leopard | *Panthera uncia* | - |
| **24** | Clouded Leopard | *Neofelis nebulosa* | - |
| **25** | Jaguarundi | *Herpailurus yagouaroundi* | - |
| **26** | Black footed cat | *Felis nigripes* | - |
| **27** | Fishing cat | *Prionailurus viverrinus* | - |
| **28** | Leopard cat | *Prionailurus bengalensis* | - |
| **29** | Lynx | *Lynx canadensis* | XM_030301258.1 |
| **30** | Striped hyena | *Hyaena hyaena* | XM_039243533.1 |
| **31** | Meerkat | *Suricata suricatta* | XM_029949927.1 |

**-** : nucleotide sequences obtained through customized local implementation of BLAST.

#### Supplementary table S4: *Zp4* accession numbers

| | Species | | Accession number |
|---|---|---|---|
| **1** | Dog | *Canis lupus familiaris* | XM_038487243.1 |
| **2** | Dingo | *Canis lupus dingo* | XM_025448645.2 |
| **3** | Red fox | *Vulpes vulpes* | XM_026008664.1 |
| **4** | Grizzly bear | *Ursus arctos horribilis* | XM_026503989.2 |
| **5** | Polar bear | *Ursus maritimus* | XM_040624263.1 |
| **6** | American black bear | *Ursus americanus* | XM_045802298.1 |
| **7** | Panda | *Ailuropoda melanoleuca* | XM_019804231.2 |
| **8** | American mink | *Neovison vison* | XM_044236554.1 |
| **9** | Sea otter | *Enhydra lutris* | XM_022500198.1 |
| **10** | Ferret | *Mustela putorius furo* | XM_045087341.1 |
| **11** | Pacific walrus | *Odobenus rosmarus* | XM_004411337.1 |
| **12** | Steller sea lion | *Eumetopias jubatus* | XM_028104607.1 |
| **13** | California sea lion | *Zalophus californianus* | XM_027596652.1 |
| **14** | Hawaiian monk seal | *Neomonachus schauinslandi* | XM_021696173.2 |
| **15** | Weddell seal | *Leptonychotes weddellii* | XM_006732888.1 |
| **16** | Meerkat | *Suricata suricatta* | XM_029915014.1 |
| **17** | Cat | *Felis catus* | NM_001009260.1 |
| **18** | Cheetah | *Acinonyx jubatus* | XM_015085463.2 |
| **19** | Puma | *Puma concolor* | XM_025915060.1 |
| **20** | Canada lynx | *Lynx canadensis* | XM_030335447.1 |
| **21** | Tiger | *Panthera tigris* | XM_007086048.2 |
| **22** | Leopard | *Panthera pardus* | XM_019419088.1 |
| **23** | Jaguar | *Panthera onca* | - |
| **24** | Lion | *Panthera leo* | - |
| **25** | Snow leopard | *Panthera uncia* | - |
| **26** | Clouded Leopard | *Neofelis nebulosa* | - |
| **27** | Jaguarundi | *Herpailurus yagouaroundi* | XM_040465019.1 |
| **28** | Black footed cat | *Felis nigripes* | - |
| **29** | Fishing cat | *Prionailurus viverrinus* | XM_047826042.1 |
| **30** | Leopard cat | *Prionailurus bengalensis* | XM_043598162.1 |
| **31** | Striped hyena | *Hyaena hyaena* | XM_039254353.1 |

**-** : nucleotide sequences obtained through customized local implementation of BLAST.

**Supplementary table S5: *Zp1* accession numbers**

| | Species | | Accession number |
|---|---|---|---|
| **1** | Ferret | *Mustela putorius furo* | XM_004770464.3 |
| **2** | American mink | *Neovison vison* | XM_044261161.1 |
| **3** | Sea otter | *Enhydra lutris* | XM_022507697.1 |
| **4** | Polar bear | *Ursus maritimus* | XM_040639172.1 |
| **5** | Panda | *Ailuropoda melanoleuca* | XM_034645141.1 |
| **6** | American black bear | *Ursus americanus* | XM_045786785.1 |
| **7** | Grizzly bear | *Ursus arctos horribilis* | XM_026487758.2 |
| **8** | Steller sea lion | *Eumetopias jubatus* | XM_028118148.1 |
| **9** | Hawaiian monk seal | *Neomonachus schauinslandi* | XM_021689794.1 |
| **10** | Pacific walrus | *Odobenus rosmarus* | XM_012565789.1 |
| **11** | California sea lion | *Zalophus californianus* | XM_027580557.2 |
| **12** | Northern fur seal | *Callorhinus ursinus* | XM_025878351.1 |
| **13** | Leopard | *Panthera pardus* | XM_019414554.1 |
| **14** | Jaguar | *Panthera onca* | - |
| **15** | Tiger | *Panthera tigris* | XM_007092010.3 |
| **16** | Cheetah | *Acinonyx jubatus* | XM_027045024.1 |
| **17** | Cat | *Felis catus* | NM_001284435.1 |
| **18** | Lion | *Panthera leo* | XM_042904576.1 |
| **19** | Snow leopard | *Panthera uncia* | XM_049645529.1 |
| **20** | Clouded Leopard | *Neofelis nebulosa* | - |
| **21** | Jaguarundi | *Herpailurus yagouaroundi* | XM_040495006.1 |
| **22** | Black footed cat | *Felis nigripes* | - |
| **23** | Fishing cat | *Prionailurus viverrinus* | XM_047848546.1 |
| **24** | Leopard cat | *Prionailurus bengalensis* | XM_043581266.1 |
| **25** | Lynx | *Lynx canadensis* | XM_030331976.1 |
| **26** | Cougar | *Puma concolor* | XM_025915635.1 |
| **27** | Striped hyena | *Hyaena hyaena* | XM_039253540.1 |
| **28** | Meerkat | *Suricata suricatta* | XM_029915453.1 |

**-** : nucleotide sequences obtained through customized local implementation of BLAST.

#### Supplementary table S6: *Izumo1r* accession numbers

| Species | | | Accession number |
|---|---|---|---|
| **1** | Red fox | *Vulpes vulpes* | XM_025993322.1 |
| **2** | Dog | *Canis lupus familiaris* | XM_038429635.1 |
| **3** | Dingo | *Canis lupus dingo* | XM_025419176.2 |
| **4** | Grizzly bear | *Ursus arctos horribilis* | XM_044386536.1 |
| **5** | Panda | *Ailuropoda melanoleuca* | XM_002930414.3 |
| **6** | Polar bear | *Ursus maritimus* | XM_040630688.1 |
| **7** | American mink | *Neovison vison* | XM_044260702.1 |
| **8** | Sea otter | *Enhydra lutris* | XM_022514185.1 |
| **9** | Ferret | *Mustela putorius furo* | XM_004754509.3 |
| **10** | Hawaiian monk seal | *Neomonachus schauinslandi* | XM_021679101.1 |
| **11** | Weddell seal | *Leptonychotes weddellii* | XM_031036823.1 |
| **12** | Pacific walrus | *Odobenus rosmarus* | XM_004410756.1 |
| **13** | California sea lion | *Zalophus californianus* | XM_027580383.2 |
| **14** | Steller sea lion | *Eumetopias jubatus* | XM_028119634.1 |
| **15** | Northern fur seal | *Callorhinus ursinus* | XM_025893961.1 |
| **16** | Jaguar | *Panthera onca* | - |
| **17** | Leopard | *Panthera pardus* | XM_019450101.1 |
| **18** | Tiger | *Panthera tigris* | XM_042958585.1 |
| **19** | Cheetah | *Acinonyx jubatus* | XM_027039662.1 |
| **20** | Cat | *Felis catus* | XM_023238987.2 |
| **21** | Lion | *Panthera leo* | - |
| **22** | Snow leopard | *Panthera uncia* | - |
| **23** | Clouded Leopard | *Neofelis nebulosa* | - |
| **24** | Jaguarundi | *Herpailurus yagouaroundi* | - |
| **25** | Black footed cat | *Felis nigripes* | - |
| **26** | Fishing cat | *Prionailurus viverrinus* | - |
| **27** | Leopard cat | *Prionailurus bengalensis* | - |
| **28** | Lynx | *Lynx canadensis* | |
| **29** | Puma | *Puma concolor* | - |
| **30** | Striped hyena | *Hyaena hyaena* | XM_039254982.1 |
| **31** | Meerkat | *Suricata suricatta* | - |

**-** : nucleotide sequences obtained through customized local implementation of BLAST.

#### Supplementary table S7: *Izumo1* accession numbers

| | Species | | Accession number |
|---|---|---|---|
| **1** | Red fox | *Vulpes vulpes* | XM_025993322.1 |
| **2** | Dog | *Canis lupus familiaris* | XM_038429635.1 |
| **3** | Dingo | *Canis lupus dingo* | XM_025419176.2 |
| **4** | Grizzly bear | *Ursus arctos horribilis* | XM_044386536.1 |
| **5** | American black bear | *Ursus americanus* | XM_040634272.1 |
| **6** | Panda | *Ailuropoda melanoleuca* | XM_002930414.3 |
| **7** | Polar bear | *Ursus maritimus* | XM_040630688.1 |
| **8** | Sea otter | *Enhydra lutris* | XM_022514185.1 |
| **9** | American mink | *Neovison vison* | XM_044260702.1 |
| **10** | Ferret | *Mustela putorius furo* | XM_004754509.3 |
| **11** | Ermin | *Mustela erminea* | |
| **12** | Northern fur seal | *Callorhinus ursinus* | XM_025893961.1 |
| **13** | Steller sea lion | *Eumetopias jubatus* | XM_028119634.1 |
| **14** | Hawaiian monk seal | *Neomonachus schauinslandi* | XM_021679101.1 |
| **15** | Weddell seal | *Leptonychotes weddellii* | XM_031036823.1 |
| **16** | Pacific walrus | *Odobenus rosmarus* | XM_004410756.1 |
| **17** | California sea lion | *Zalophus californianus* | XM_027580383.2 |
| **18** | Jaguar | *Panthera onca* | - |
| **19** | Leopard | *Panthera pardus* | XM_019450101.1 |
| **20** | Tiger | *Panthera tigris* | XM_042958585.1 |
| **21** | Cheetah | *Acinonyx jubatus* | XM_027039662.1 |
| **22** | Cat | *Felis catus* | XM_023238987.2 |
| **23** | Lion | *Panthera leo* | XM_042917797.1 |
| **24** | Snow leopard | *Panthera uncia* | - |
| **25** | Clouded Leopard | *Neofelis nebulosa* | - |
| **26** | Jaguarundi | *Herpailurus yagouaroundi* | - |
| **27** | Black footed cat | *Felis nigripes* | - |
| **28** | Fishing cat | *Prionailurus viverrinus* | - |
| **29** | Leopard cat | *Prionailurus bengalensis* | XM_043600374.1 |
| **30** | Lynx | *Lynx canadensis* | XM_030298692.1 |
| **31** | Cougar | *Puma concolor* | XM_025914312.1 |
| **32** | Striped hyena | *Hyaena hyaena* | XM_039247704.1 |
| **33** | Meerkat | *Suricata suricatta* | XM_029924449.1 |

**-** : nucleotide sequences obtained through customized local implementation of BLAST.

**Supplementary table S8: pairwise sequence identities report for *Carnivora* in each studied protein**

| | | | ZP2 | ZP3 | ZP4 | ZP1 | JUNO | IZUMO1 |
|---|---|---|---|---|---|---|---|---|
| *Carnivora* | Identity | Mean | 85.22 | 86.18 | 84.16 | 86.29 | 85.95 | 67.26 |
| | | Minimun | 72.75 | 75.96 | 73.23 | 69.20 | 59.47 | 45.66 |
| | | Maximun | 99.86 | 100.00 | 99.64 | 99.67 | 100.00 | 100.00 |
| | | Standar deviation | 7.70 | 6.87 | 8.61 | 9.37 | 8.48 | 17.87 |
| | Number of species | | 31.00 | 31.00 | 31.00 | 28.00 | 31.00 | 33.00 |

Mean, minimum, maximum, and standard deviation of sequence identity comparisons were determined for each protein using comprehensive alignments that encompassed all carnivore species incorporated in the analysis

| source | sum of squares SS | degrees of freedom ν | mean square MS | p-value |
|---|---|---|---|---|
| treatment | 1,366,698,103 | 5 | 273,339,621 | 1.11E-12 |
| error | 3,193,072,470 | 2760 | 1,156,910 | |
| total | 4,559,770,572 | 2765 | | |

One-way ANOVA of k=6 independent treatments

| treatments pair | Scheffé T-statistic | Scheffé p-value | Scheffé inferfence |
|---|---|---|---|
| IZUMO1 vs JUN( | 273,161 | 1.11E-12 | ** p<0.01 |
| IZUMO1 vs ZP3 | 262,520 | 1.11E-12 | ** p<0.01 |
| IZUMO1 vs ZP2 | 276,582 | 1.11E-12 | ** p<0.01 |
| IZUMO1 vs ZP4 | 247,032 | 1.11E-12 | ** p<0.01 |
| IZUMO1 vs ZP1 | 212,818 | 1.11E-12 | ** p<0.01 |
| JUNO vs ZP3 | 1.0318 | 9,571,473 | insignificant |
| JUNO vs ZP2 | 3,317 | 9,997,944 | insignificant |
| JUNO vs ZP4 | 2.5337 | 2,678,294 | insignificant |
| JUNO vs ZP1 | 4.3800 | 18,191 | ** p<0.01 |
| ZP3 vs ZP2 | 1.3635 | 8,682,066 | insignificant |
| ZP3 vs ZP4 | 1.5019 | 8,127,101 | insignificant |
| ZP3 vs ZP1 | 3.4028 | 413,181 | * p<0.05 |
| ZP2 vs ZP4 | 2.8654 | 1,453,984 | insignificant |
| ZP2 vs ZP1 | 4.6941 | 5,331 | ** p<0.01 |
| ZP4 vs ZP1 | 1.9806 | 5,607,054 | insignificant |

Scheffé results

| | |
|---|---|
| **p value < 0.05** | **Significativo** |
| **p value > 0.05** | **No significativo** |

**Supplementary table S9: pairwise sequence identities report for taxonimic groups of interest in each studied protein and negative control**

| | | | Caniformia | Feliformia | Panterinae |
|---|---|---|---|---|---|
| ZP3 | Identity | Mean | 88.32 | 95.49 | 98.75 |
| | | Minimun | 80.98 | 86.55 | 98.11 |
| | | Maximun | 100.00 | 100.00 | 100.00 |
| | | Standar deviation | 4.61 | 4.53 | 0.48 |
| | Number of species | | 16.00 | 15.00 | 6.00 |
| | p-value T-Test | | 0.0002569 *** | | |
| ZP2 | Identity | Mean | 85.62 | 94.92 | 98.52 |
| | | Minimun | 75.13 | 85.49 | 96.51 |
| | | Maximun | 100.00 | 100.00 | 99.58 |
| | | Standar deviation | 6.63 | 4.46 | 1.11 |
| | Number of species | | 15.00 | 16.00 | 7.00 |
| | p-value T-Test | | 0.0001261 *** | | |
| ZP4 | Identity | Mean | 89.22 | 93.80 | 98.31 |
| | | Minimun | 81.08 | 79.75 | 97.18 |
| | | Maximun | 99.64 | 99.47 | 99.47 |
| | | Standar deviation | 5.05 | 6.50 | 0.66 |
| | Number of species | | 15.00 | 16.00 | 6.00 |
| | p-value T-Test | | 0.03649 * | | |
| ZP1 | Identity | Mean | 83.29 | 93.74 | 98.62 |
| | | Minimun | 72.10 | 80.00 | 97.98 |
| | | Maximun | 99.35 | 99.67 | 99.26 |
| | | Standar deviation | 6.91 | 6.21 | 0.40 |
| | Number of species | | 12.00 | 16.00 | 6.00 |
| | p-value T-Test | | 0.0004252 *** | | |
| JUNO | Identity | Mean | 84.60 | 96.21 | 99.10 |
| | | Minimun | 67.07 | 87.80 | 98.40 |
| | | Maximun | 100.00 | 100.00 | 100.00 |
| | | Standar deviation | 8.23 | 4.22 | 0.42 |
| | Number of species | | 15.00 | 16.00 | 6.00 |
| | p-value T-Test | | 0.00008105 *** | | |
| IZUMO1 | Identity | Mean | 71.65 | 91.79 | 97.79 |
| | | Minimun | 48.33 | 73.05 | 95.95 |
| | | Maximun | 100.00 | 99.13 | 98.84 |
| | | Standar deviation | 16.17 | 8.03 | 0.92 |
| | Number of species | | 17.00 | 16.00 | 6.00 |
| | p-value T-Test | | 0.0001276 *** | | |
| ACTN1 | Identity | Mean | 99.65 | 99.84 | 99.93 |
| | | Minimun | 95.74 | 99.01 | 99.78 |
| | | Maximun | 100.00 | 100.00 | 100.00 |
| | | Standar deviation | 0.48 | 0.26 | 0.07 |
| | Number of species | | 15.00 | 16.00 | 6.00 |
| | p-value T-Test | | 0.2058 NS | | |

Identity calculation were performed using Sequence Identity And Similarity online tool (SIAS – http://imed.med.ucm.es/Tools/sias.html).
Caniformes vs Feliformes mean identity comparison for each protein were performed through a two sample T-Test (Welch's T-test).

| | |
|---|---|
| **p value < 0.05** | **Significativo** |
| **p value > 0.05** | **No significativo** |

**Supplementary table S10: branch-site positive selection results for *Zp2***

| *Zp2* | | | | | | |
|---|---|---|---|---|---|---|
| | **Log Likelihood** | **LRT** | **p** | **Parameters Estimation** | | |
| **Caniformia** | | | | | | |
| Basal branch | | | | | | |
| Alternative hypothesis | -9,007.4560 | | | p0 = 0.62671 | p1 = 0.37329 | p2 = 0.0000 |
| Null hypothesis | -9,007.4560 | 0.000 | 1.000 | ω0 = 0.17339 | ω1 = 1.0000 | ω2 = 1.0000 |
| Subtree analysis | | | | | | |
| Alternative hypothesis | -9,004.94 | | | p0 = 0.62383 | p1 = 0.27776 | p2 = 0.09841 |
| Null hypothesis | -9,005.39 | 0.892 | 0.345 | ω0 = 0.17177 | ω1 = 1.0000 | ω2 = 1.49101 |
| **Feliformia** | | | | | | |
| Basal branch | | | | | | |
| Alternative hypothesis | -9,007.46 | | | p0 = 0.62671 | p1 = 0.37329 | p2 = 0.0000 |
| Null hypothesis | -9,007.46 | 0.000 | 1.000 | ω0 = 0.17399 | ω1 = 1.0000 | ω2 = 1.24902 |
| Subtree analysis | | | | | | |
| Alternative hypothesis | -8,996.00 | | | p0 = 0.62485 | p1 = 0.37244 | p2 = 0.00271 |
| Null hypothesis | -9,007.37 | 22.742 | <0.00001 | ω0 = 0.17615 | ω1 = 1.0000 | ω2 = 16.34637 |
| **Pantherinae** | | | | | | |
| Basal branch | | | | | | |
| Alternative hypothesis | -9,007.46 | | | p0 = 0.62671 | p1 = 0.37329 | p2 = 0.0000 |
| Null hypothesis | -9,007.46 | 0.000 | 1.000 | ω0 = 0.17399 | ω1 = 1.0000 | ω2 = 1.0000 |
| Subtree analysis | | | | | | |
| Alternative hypothesis | -8,999.21 | | | p0 = 0.62052 | p1 = 0.3779 | p2 = 0.00151 |
| Null hypothesis | -9,007.46 | 16.503 | <0.00001 | ω0 = 0.17399 | ω1 = 1.0000 | ω2 = 136.63784 |

LRT: statistical value of the likelihood ratio test; p0-2 and ω0-2 proportion of sites and dN/dS values estimated for each site class in the selection model (alternative hypothesis) respecively.

| | |
|---|---|
| **p value < 0.05** | **Significativo** |
| **p value > 0.05** | **No significativo** |

**Supplementary table S11: branch-site positive selection results for *Zp3***

| *Zp3* | | | | | | |
|---|---|---|---|---|---|---|
| | **Log Likelihood** | **LRT** | **p** | **Parameters Estimation** | | |
| **Caniformia** | | | | | | |
| Basal branch | | | | | | |
| Alternative hypothesis | -5,801.7500 | | | p0 = 0.74747 | p1 = 0.23110 | p2 = 0.02143 |
| Null hypothesis | -5,801.7500 | 0.000 | 1.000 | ω0 = 0.09830 | ω1 = 1.0000 | ω2 = 1.0000 |
| Subtree analysis | | | | | | |
| Alternative hypothesis | -5,797.85 | | | p0 = 0.76549 | p1 = 0.00296 | p2 = 0.01583 |
| Null hypothesis | -5,801.32 | 6.945 | 0.008 | ω0 = 0.10315 | ω1 = 1.0000 | ω2 = 10.40742 |
| **Feliformia** | | | | | | |
| Basal branch | | | | | | |
| Alternative hypothesis | -5,800.36 | | | p0 = 0.76279 | p1 = 0.22064 | p2 = 0.01658 |
| Null hypothesis | -5,800.36 | 0.000 | 1.000 | ω0 = 0.10377 | ω1 = 1.0000 | ω2 = 11.96120 |
| Subtree analysis | | | | | | |
| Alternative hypothesis | -5,800.14 | | | p0 = 0.72628 | p1 = 0.20732 | p2 = 0.06639 |
| Null hypothesis | -5,800.14 | 0.000 | 1.000 | ω0 = 0..08858 | ω1 = 1.0000 | ω2 = 1.0000 |
| **Pantherinae** | | | | | | |
| Basal branch | | | | | | |
| Alternative hypothesis | -5,801.84 | | | p0 = 0.76438 | p1 = 0.23562 | p2 = 0.0000 |
| Null hypothesis | -5,801.84 | 0.000 | 1.000 | ω0 = 0.10067 | ω1 = 1.0000 | ω2 = 1.0000 |
| Subtree analysis | | | | | | |
| Alternative hypothesis | -5,801.84 | | | p0 = 0.76438 | p1 = 0.23562 | p2 = 0.0000 |
| Null hypothesis | -5,801.84 | 0.000 | 1.000 | ω0 = 0.10067 | ω1 = 1.0000 | ω2 = 1.0000 |

LRT: statistical value of the likelihood ratio test; p0-2 and ω0-2 proportion of sites and dN/dS values estimated for each site class in the selection model (alternative hypothesis) respecively.

| | |
|---|---|
| **p value < 0.05** | **Significativo** |
| **p value > 0.05** | **No significativo** |

**Supplementary table S12: branch-site positive selection results for *Zp4***

| *Zp4* | | | | | | |
|---|---|---|---|---|---|---|
| | **Log Likelihood** | **LRT** | **p** | **Parameters Estimation** | | |
| **Caniformia** | | | | | | |
| Basal branch | | | | | | |
| Alternative hypothesis | -7,808.1300 | | | p0 = 0.62426 | p1 = 0.34881 | p2 = 0.02692 |
| Null hypothesis | -7,810.2800 | 4.286 | 0.038 | ω0 = 0.14704 | ω1 = 1.0000 | ω2 = 7.87390 |
| Subtree analysis | | | | | | |
| Alternative hypothesis | -7,809.88 | | | p0 = 0.62599 | p1 = 0.30621 | p2 = 0.067780 |
| Null hypothesis | -7,810.24 | 0.722 | 0.395 | ω0 = 0.14794 | ω1 = 1.0000 | ω2 = 1.44150 |
| **Feliformia** | | | | | | |
| Basal branch | | | | | | |
| Alternative hypothesis | -7,811.07 | | | p0 = 0.62803 | p1 = 0.37197 | p2 = 0.0000 |
| Null hypothesis | -7,811.07 | 0.000 | 1.000 | ω0 = 0.14899 | ω1 = 1.0000 | ω2 = 1.0000 |
| Subtree analysis | | | | | | |
| Alternative hypothesis | -7,809.59 | | | p0 = 0.56455 | p1 = 0.31209 | p2 = 0.12335 |
| Null hypothesis | -7,809.59 | 0.000 | 1.000 | ω0 = 0.12433 | ω1 = 1.0000 | ω2 = 1.0000 |
| **Pantherinae** | | | | | | |
| Basal branch | | | | | | |
| Alternative hypothesis | -7,811.01 | | | p0 = 0.62020 | p1 = 0.36693 | p2 = 0.01278 |
| Null hypothesis | -7,811.05 | 0.086 | 0.770 | ω0 = 0.14857 | ω1 = 1.0000 | ω2 = 10.51650 |
| Subtree analysis | | | | | | |
| Alternative hypothesis | -7,809.25 | | | p0 = 0.61997 | p1 = 0.35840 | p2 = 0.02163 |
| Null hypothesis | -7,810.12 | 1.724 | 0.188 | ω0 = 0.14470 | ω1 = 1.0000 | ω2 = 6.41271 |

LRT: statistical value of the likelihood ratio test; p0-2 and ω0-2 proportion of sites and dN/dS values estimated for each site class in the selection model (alternative hypothesis) respecively.

| | |
|---|---|
| **p value < 0.05** | **Significativo** |
| **p value > 0.05** | **No significativo** |

**Supplementary table S13: branch-site positive selection results for *Zp1***

| *Zp1* | | | | | | |
|---|---|---|---|---|---|---|
| | **Log Likelihood** | **LRT** | **p** | **Parameters Estimation** | | |
| **Caniformia** | | | | | | |
| Basal branch | | | | | | |
| Alternative hypothesis | -9,093.6580 | 0.282 | 0.282 | p0 = 0.70444 | p1 = 0.29069 | p2 = 0.0487 |
| Null hypothesis | -9,093.7990 | | | ω0 = 0.17603 | ω1 = 1.0000 | ω2 = 6.18812 |
| Subtree analysis | | | | | | |
| Alternative hypothesis | -9,081.14 | 0.000 | 0.000 | p0 = 0.59213 | p1 = 0.15887 | p2 = 0.024899 |
| Null hypothesis | -9,081.14 | | | ω0 = 0.13110 | ω1 = 1.0000 | ω2 = 1.0000 |
| **Feliformia** | | | | | | |
| Basal branch | | | | | | |
| Alternative hypothesis | -9,092.00 | 1.834 | 1.834 | p0 = 0.69704 | p1 = 0.27990 | p2 = 0.02306 |
| Null hypothesis | -9,092.91 | | | ω0 = 0.17351 | ω1 = 1.0000 | ω2 = 4.32064 |
| Subtree analysis | | | | | | |
| Alternative hypothesis | -9,093.46 | 0.213 | 0.645 | p0 = 0.70018 | p1 = 0.28015 | p2 = 0.01967 |
| Null hypothesis | -9,093.57 | | | ω0 = 0.17680 | ω1 = 1.0000 | ω2 = 1.61741 |
| **Pantherinae** | | | | | | |
| Basal branch | | | | | | |
| Alternative hypothesis | -9,093.80 | 0.000 | 1.000 | p0 = 0.70644 | p1 = 0.29356 | p2 = 0.0000 |
| Null hypothesis | -9,093.80 | | | ω0 = 0.17680 | ω1 = 1.0000 | ω2 = 1.0000 |
| Subtree analysis | | | | | | |
| Alternative hypothesis | -9,093.80 | 0.000 | 1.000 | p0 = 0.70644 | p1 = 0.29356 | p2 = 0.0000 |
| Null hypothesis | -9,093.80 | | | ω0 = 0.17680 | ω1 = 1.0000 | ω2 = 1.0000 |

LRT: statistical value of the likelihood ratio test; p0-2 and ω0-2 proportion of sites and dN/dS values estimated for each site class in the selection model (alternative hypothesis) respecively.

| | |
|---|---|
| **p value < 0.05** | **Significativo** |
| **p value > 0.05** | **No significativo** |

# ZP2

A

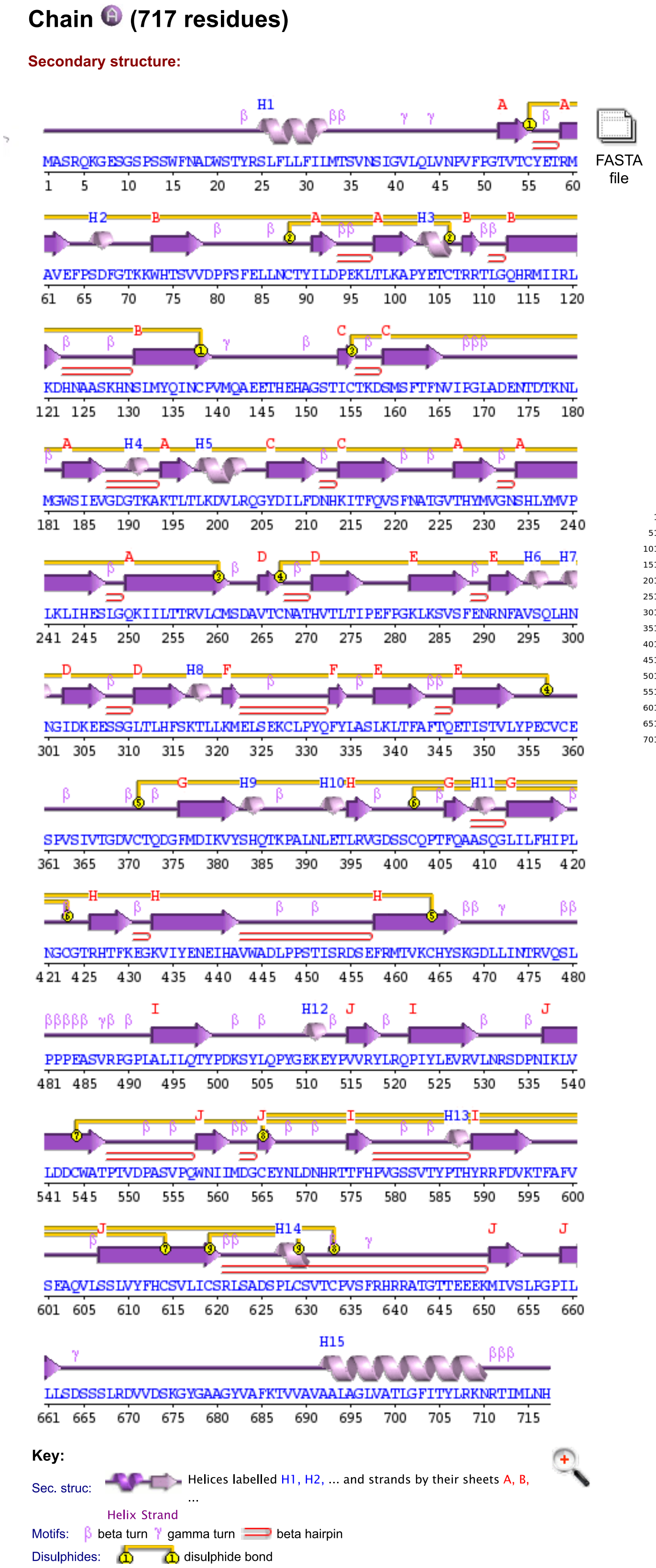


B

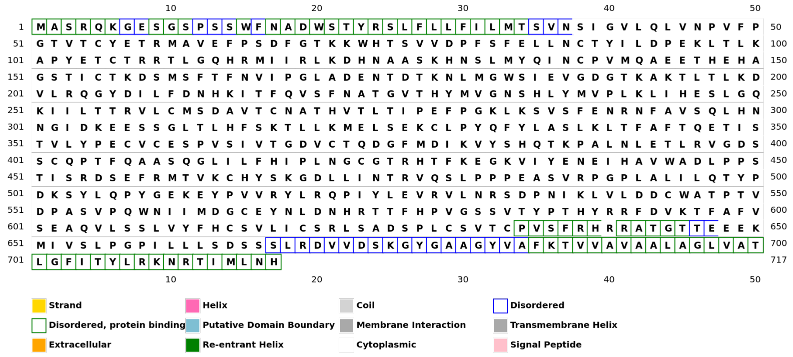


C

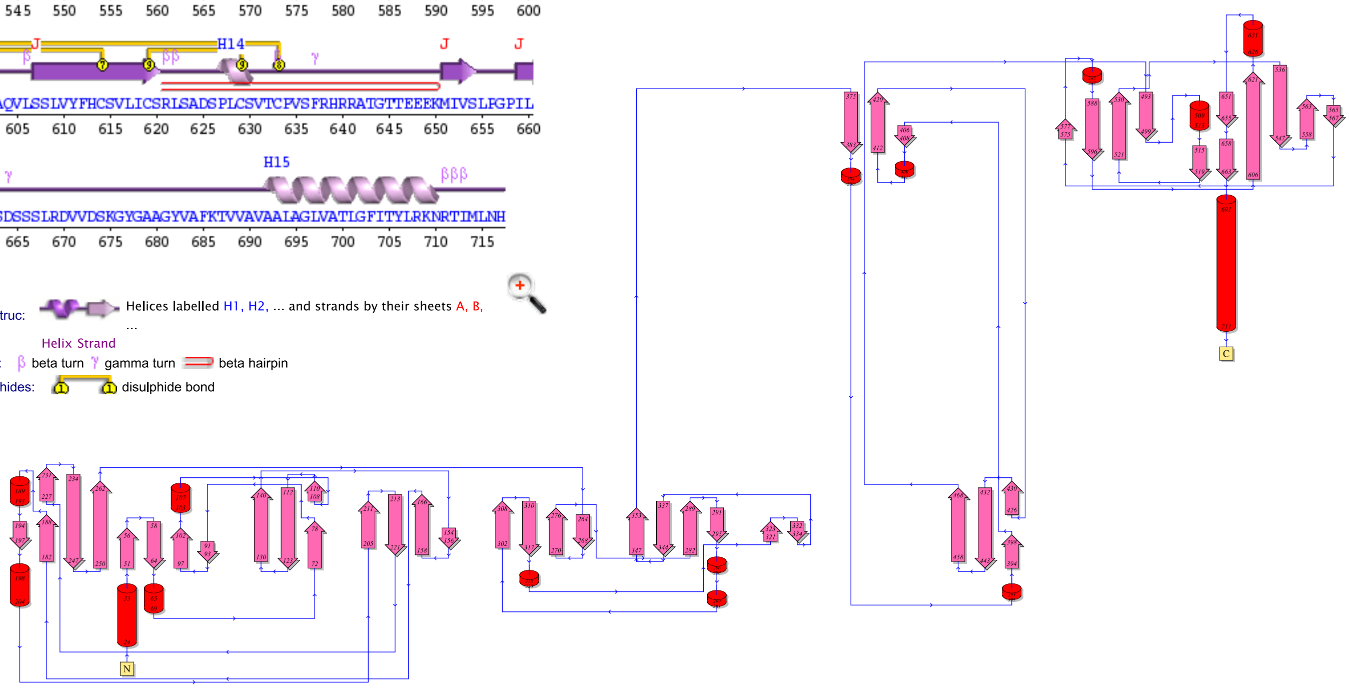


*A. Secondary structure of ZP2 protein*
*B. DISOPRED plot*
*C. Topology of ZP2 protein*

# ZP2



| Species | ZP-N2 (227–235; site 231) | ZP-N3 (320–328; site 324) |
|---|---|---|
| **Jaguar** | THYMVGNSH | LKMELSEKC |
| *Leopard* | THYMQGNSH | LKMELSEKC |
| *South african lion* | THYMVGNSH | LKMEFSEKC |
| *Tiger* | THYMQGNSH | LKMELSEKC |
| *Clouded leopard* | THYMVGNSH | LKMEFSEKC |
| *Snow leopard* | THYMVGNSH | LKMELSEKC |
| *Cat* | THYMQGNSH | LKMEFSEKC |
| *Leopard cat* | THYMVGNSH | LKMEVSEKC |
| *Black footed cat* | THYMVGNSH | LKMEVSEKC |
| *Fishing cat* | THYMVGNSH | LKMEVSEKC |
| *Canada lynx* | THYMQGNSH | IKMEFSEKC |
| *Cheetah* | THYMQGNSH | LKMEFSEKC |
| *Puma* | THYMVGNSH | LKMEVSEKC |
| *Jaguarundi* | THYMVGNSH | LKMEFSEKC |
| *Meerkat* | THYMQGNSH | LKMKFSEKC |
| *Striped hyena* | THYMQGNSH | LKMKFSEKC |
| *Dog* | THYMQGNSH | LKMNSSEKC |
| *Dingo* | THYMQGNSH | LKMNSSEKC |
| *Red_fox* | THYMQGNSH | LKMNSSEKC |
| *American black bear* | THYAQGNSH | LKIKSSEKC |
| *Grizzly bear* | THYAQGNSH | LKIKSSEKC |
| *Panda* | THYAQGNSQ | LKIKSSEKC |
| *Polar bear* | THYAQGNSH | LKIKSSEKC |
| *California sea lion* | THYVQGNSH | LKMKSSEKC |
| *Pacific walrus* | THYVQGNSH | LKMKSSEKC |
| *Steller sea lion* | THYVQGNSH | LKMKSSEKC |
| *Wedell seal* | THYVQGNSH | LRMKSSEKC |
| *Hawaiian monk seal* | THYVQGNSH | LKMKSSEKC |
| *Sea otter* | TDYVQGNRH | LKIKSSEKC |
| *American mink* | TDYVQGNRH | LKIKSSEKC |
| *Ferret* | TDYVQGNRH | LKIKSSEKC |

*Alignment detail of the ZP2 protein to showcase positively selected sites. Diversifying positive selection in the sub-tree is highlighted by coloring amino acids in bold green lettering, while episodic positive selection on the basal branch of the clade is indicated by a green background.*

# ZP2

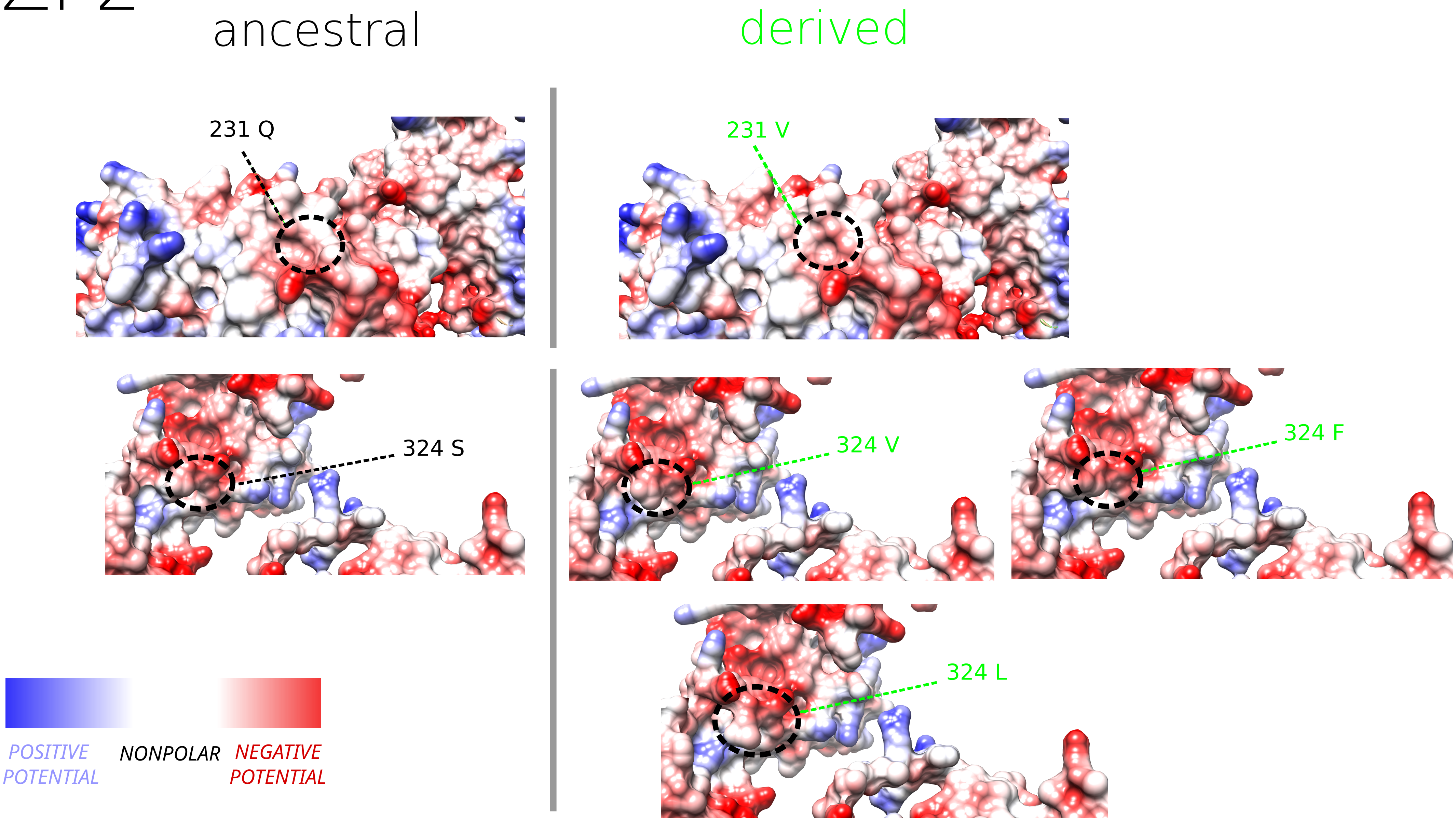


*The ancestral and derived states of positively selected amino acids depicted through Coulombic surfaces. These surfaces illustrate the electrostatic potential as governed by Coulomb's law: red indicates negative potential, blue represents positive potential, and white denotes relatively nonpolar potential.*

# ZP3

A

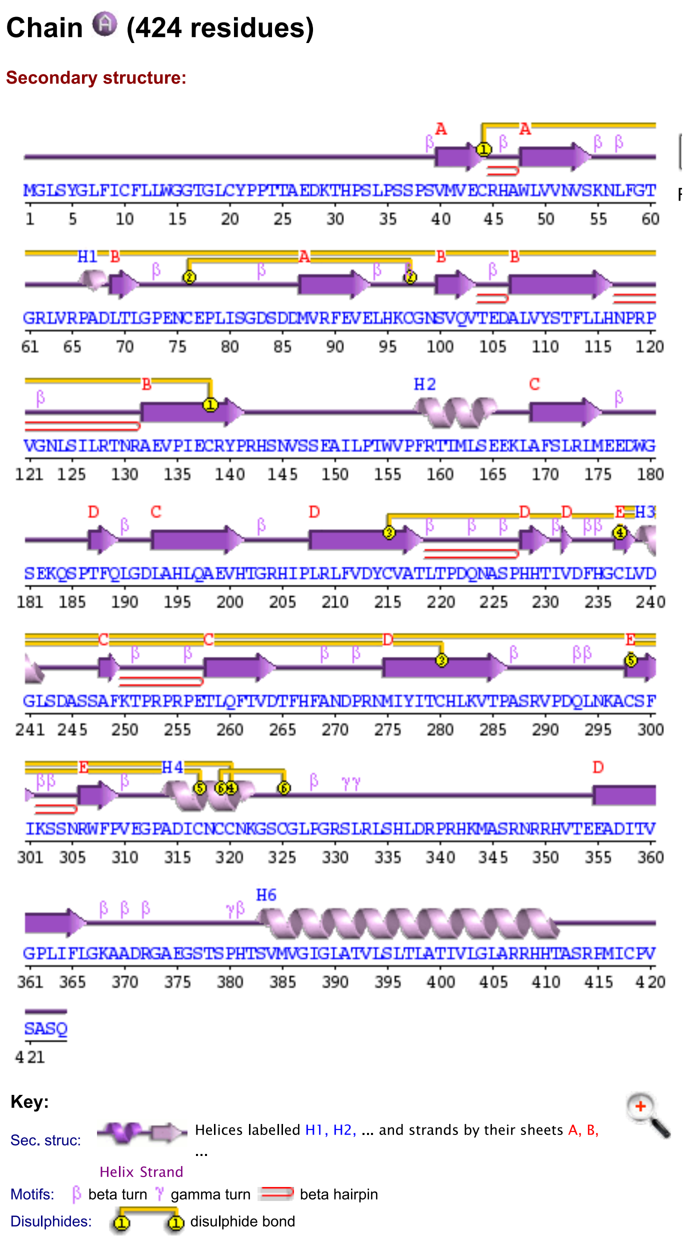


B

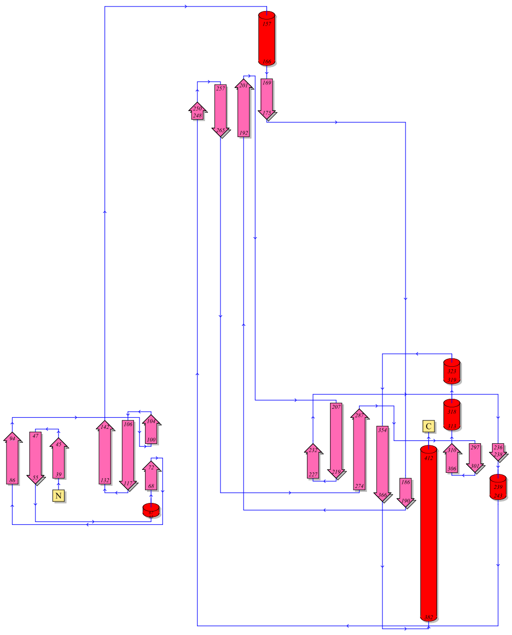

C

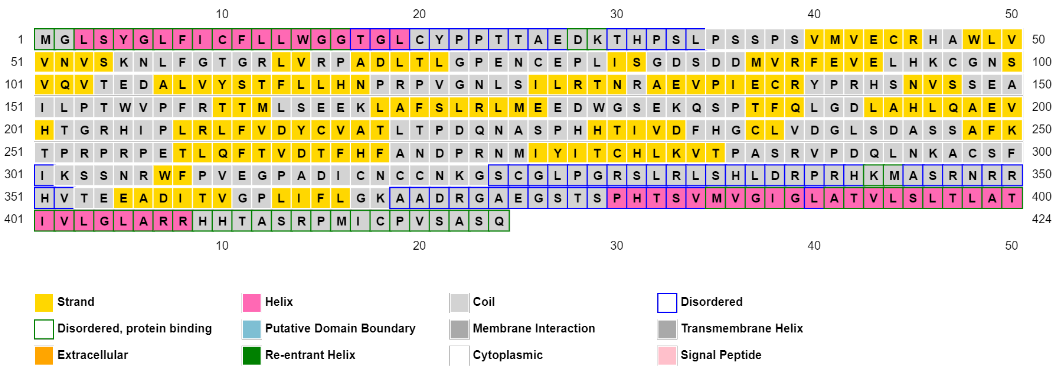


*A. Secondary structure of ZP3 protein*
*B. Topology of ZP3 protein*
*C. DISOPRED plot*

# ZP3



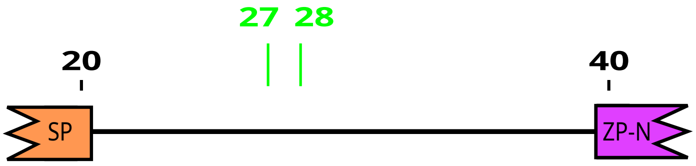


| Species | Sequence |
|---|---|
| **Jaguar** | LCYPPTTAEDKTHPSLPSSPSVM |
| *Leopard* | LCYPPTTAEDKTHPSLPSSPSVM |
| *South african lion* | LCYPPTTGEDKTHPSLPSSPSVV |
| *Tiger* | LCYPPTTAGDKTHPSLPSSPSVM |
| *Clouded leopard* | LCYPPTTGEDKTHPSLPSSPSVV |
| *Snow leopard* | LCYPPTTAGDKTHPSLPSSPSVM |
| *Cat* | LCYPPTTTEDKTHPSLPSSPSVV |
| *Leopard cat* | LCYPPTT?EDKTHPSLPSSPSVV |
| *Black footed cat* | LCYPPTTTEDKTHPSLPSSPSVV |
| *Fishing cat* | LCYPPTTTEDKTHPSLPSSPSVV |
| *Canada lynx* | LCYPPTTTEDKTHPSLPSSPSVV |
| *Cheetah* | LCYPPTTTEDKTHPSLPSSPSVV |
| *Puma* | LCYPPTTTEDKTHPSLPSSPSVV |
| *Jaguarundi* | LCYPPTTTEDKTHPSLPSSPSVV |
| *Meerkat* | LCCPQTTP-DKTHPSLPSSPPVV |
| *Striped hyena* | LCFPQTTP-DKTHPSLPSSPPVV |
| *Dog* | LCCPQTIWPTETYYPLTSRPPVM |
| *Dingo* | LCCPQTIWPTETYYPLTSRPPVM |
| *Red_fox* | LCCPQTIWPTETYYPLTSRPPVM |
| *American black bear* | LCYSQTTWWGETPYPLPSRPPVV |
| *Grizzly bear* | LCYSQTTWWGETPYPLPSRPPVV |
| *Panda* | LCYSQTTWWGETPYPLPSRPPVV |
| *Polar bear* | LCYLQTTPWRETYYSLPSRPPVV |
| *California sea lion* | LCYLQTTSQRETYYSLPPRPPVV |
| *Pacific walrus* | LCYLQTTPWRETYYSLPSRPPVV |
| *Steller sea lion* | LCYLQTTPWRETYYSLPSRPPVV |
| *Northern fur seal* | LCYPQTTSPGETYYSLPSRPPVV |
| *Wedell seal* | LCYPQTTSPGETYYSLPSRPPVV |
| *Hawaiian monk seal* | LCYSQTIWSGETNSPLPSRPPVV |
| *Sea otter* | PCCSQTIWSSETSSLLPSRPPVV |
| *American mink* | LCYPQTIWSRETSSPLPSRPPVV |
| *Ferret* | LCYSQTIWSGETSSPLPSRPPVV |
| *Ermine* | |

*Alignment detail of the ZP3 protein to showcase positively selected sites. Diversifying positive selection in the sub-tree is highlighted by coloring amino acids in bold green lettering, while episodic positive selection on the basal branch of the clade is indicated by a green background.*

# ZP3

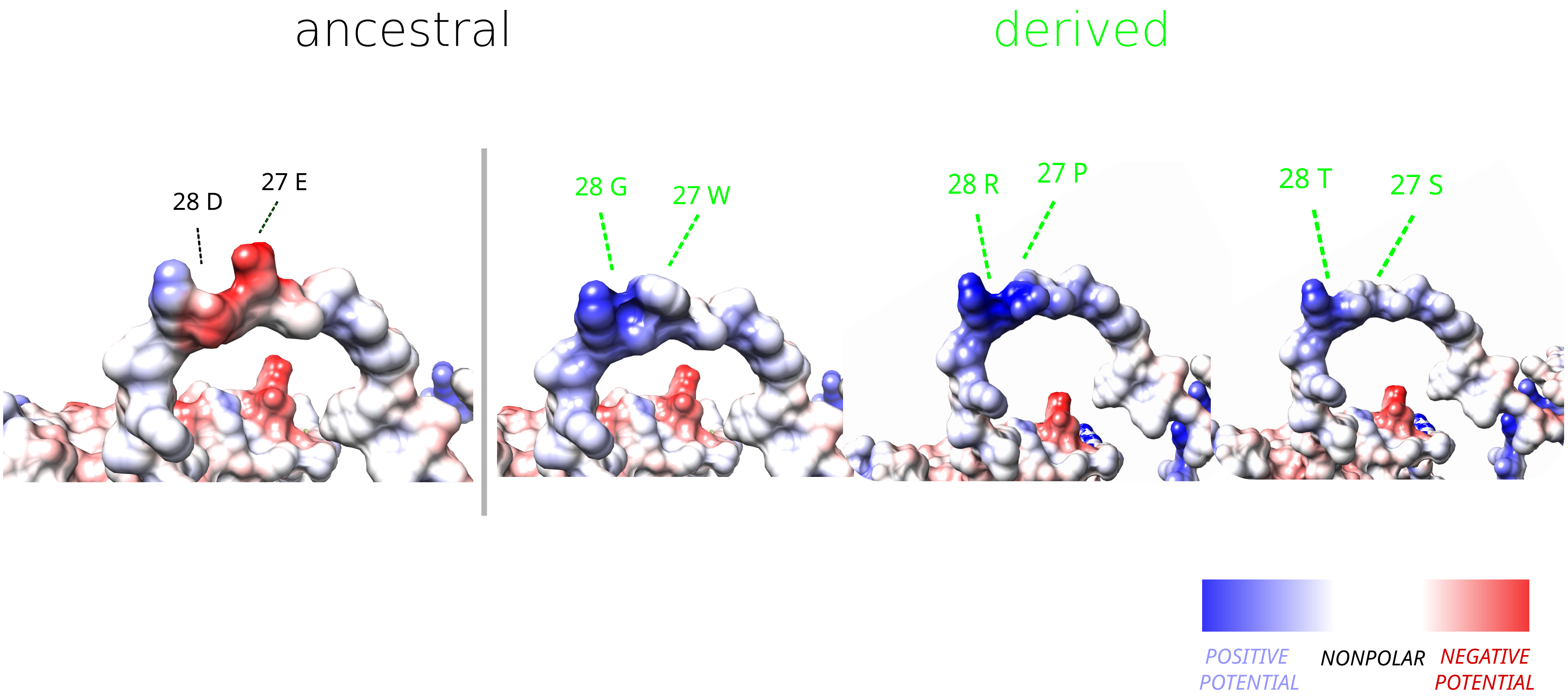


*The ancestral and derived states of positively selected amino acids depicted through Coulombic surfaces. These surfaces illustrate the electrostatic potential as governed by Coulomb's law: red indicates negative potential, blue represents positive potential, and white denotes relatively nonpolar potential.*

A

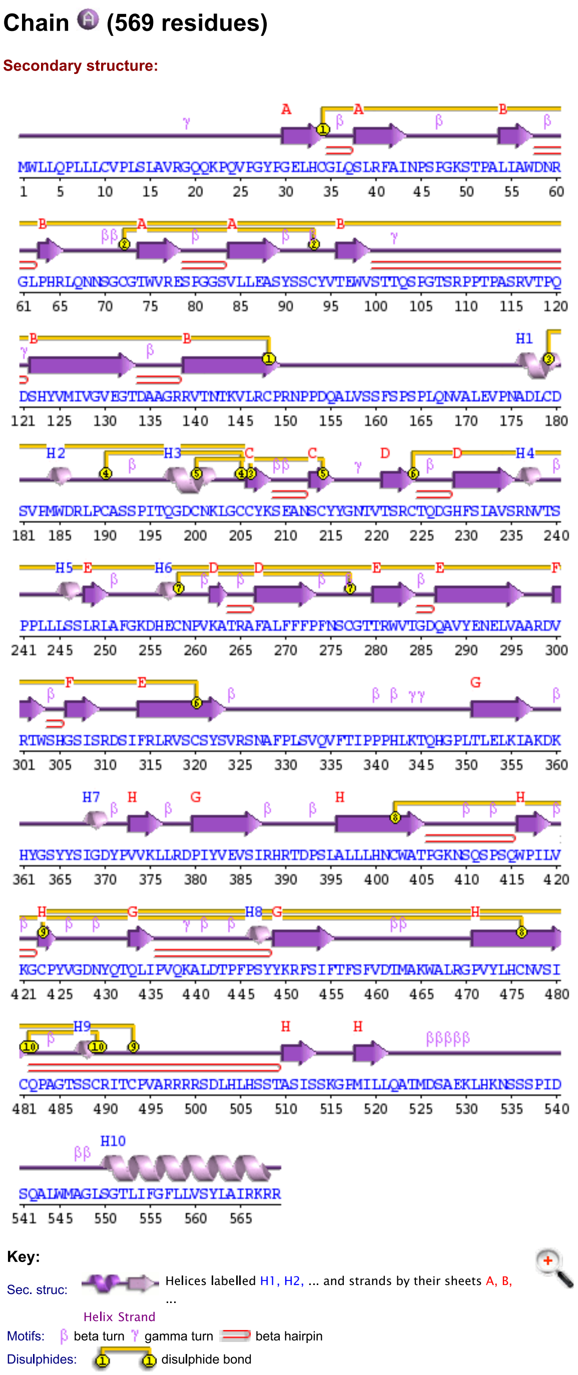


B

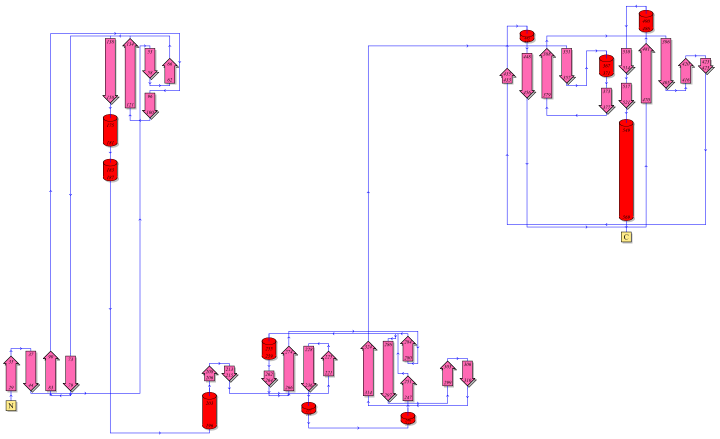

C

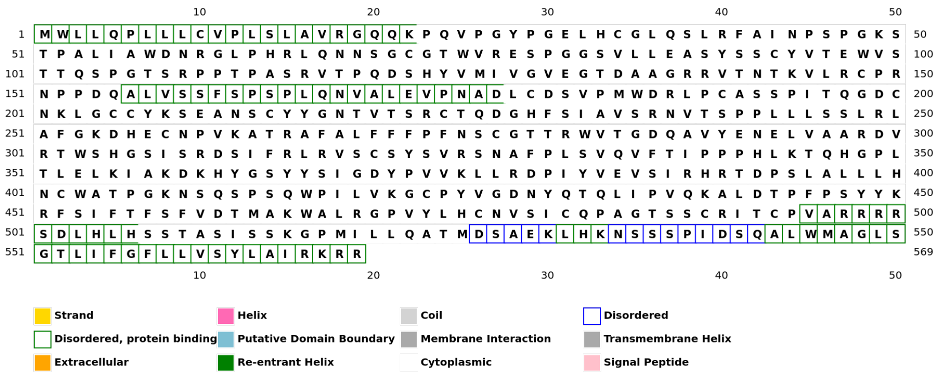


*A. Secondary structure of ZP4 protein*
*B. Topology of ZP4 protein*
*C. DISOPRED plot*

# ZP4

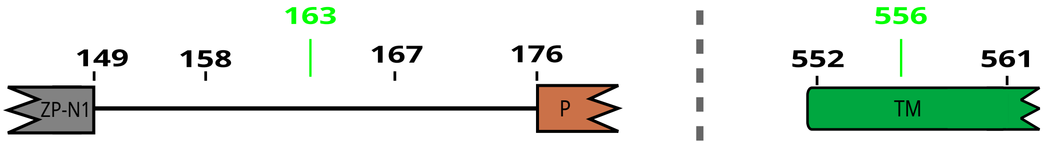


| Species | 149–176 | 552–561 |
|---|---|---|
| **Jaguar** | M W L L Q P L L L C | L I F G F L L V S Y |
| *Leopard* | M W L L Q P L L L C | L I F G F L L V S Y |
| *South african lion* | M W L L Q P L L L C | L I F G F L L V S Y |
| *Tiger* | M W L L Q P L L L C | L I F G F L L V S Y |
| *Clouded leopard* | V S S F S P S P L Q | L I F G F L L V S Y |
| *Snow leopard* | V S S F S P S P L Q | L I F G F L L V S Y |
| *Cat* | V S S L S P S P L Q | L I F G F L L V S Y |
| *Leopard cat* | V S S L S P S P L Q | L I F G F L L V S Y |
| *Black footed cat* | V S S L S P S P L Q | L I F G F L L V S Y |
| *Fishing cat* | V S S L S P S P L Q | L I F G F L L V S Y |
| *Canada lynx* | V S S S S P S P L Q | L I F G F L L V S Y |
| *Cheetah* | V S S L S P S P L Q | L I F G F L L V S Y |
| *Puma* | V S S L S P S P L Q | L I F G F L L V S Y |
| *Jaguarundi* | V S S L S P S P L Q | L I F G F L L V S Y |
| *Meerkat* | V S S V S P S P L Q | L I V G F L L V T Y |
| *Striped hyena* | V S S V S P S P L Q | L I V G F L L V S Y |
| *Dog* | L S S L S **Y** S P D Q | L I V G **A** L L V S Y |
| *Dingo* | L S S L S **Y** S P D Q | L I V G **A** L L V S Y |
| *Red_fox* | L S S L S **Y** S P D Q | L I V G **A** L L V S Y |
| *American black bear* | L S S - - **-** S P L Q | L I F G **A** L L V S Y |
| *Grizzly bear* | L S S - - **-** S P L Q | L I I G **A** L L V S Y |
| *Panda* | L S S - - **-** S P L Q | L I I G **A** L L V S Y |
| *Polar bear* | L S S - - **-** S P L Q | L I I G **A** L L V S Y |
| *California sea lion* | L S S S S **Y** S P L Q | L I I G **A** L L V S Y |
| *Pacific walrus* | - - - - - **-** - - - - | L I I G **A** L L V S Y |
| *Steller sea lion* | L S S S S **Y** S P L Q | L I I G **A** L L V S Y |
| *Wedell seal* | L S S S S **Y** L P L Q | L I I G **A** L L V S Y |
| *Hawaiian monk seal* | L S S L S **Y** S P L Q | L I I G **A** L L V S Y |
| *Sea otter* | L S S - - **Y** S P L Q | L I T G **A** L L V S Y |
| *American mink* | - - - - - **Y** S P L Q | L I I G **A** L L V S Y |
| *Ferret* | - - - - - **-** - - - - | L I I G **A** L L V S Y |

*Alignment detail of the ZP4 protein to showcase positively selected sites. Diversifying positive selection in the sub-tree is highlighted by coloring amino acids in bold green lettering, while episodic positive selection on the basal branch of the clade is indicated by a green background.*

# ZP4

*Coulombic surface*

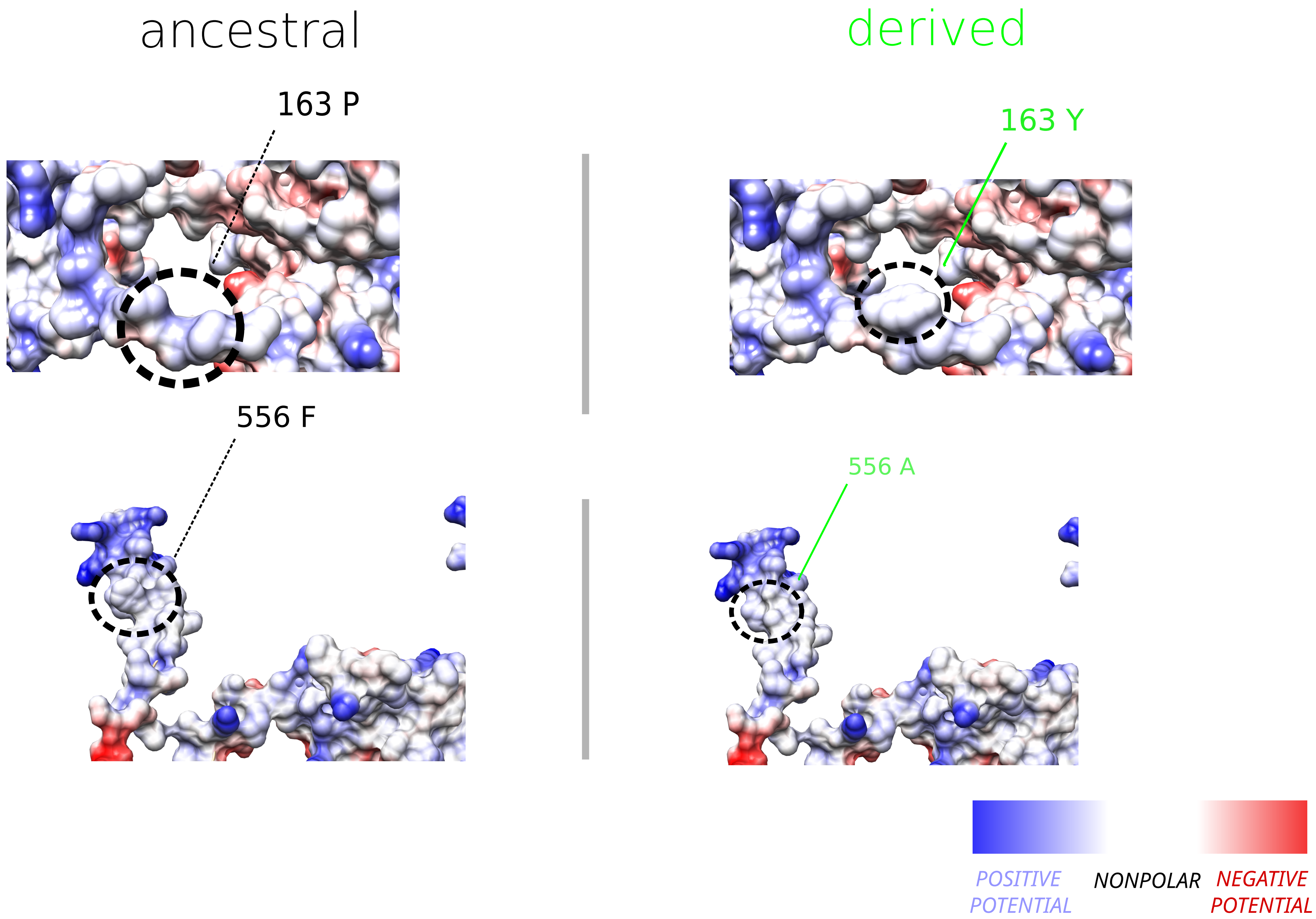


*The ancestral and derived states of positively selected amino acids depicted through Coulombic surfaces. These surfaces illustrate the electrostatic potential as governed by Coulomb's law: red indicates negative potential, blue represents positive potential, and white denotes relatively nonpolar potential.*

# ZP1

A

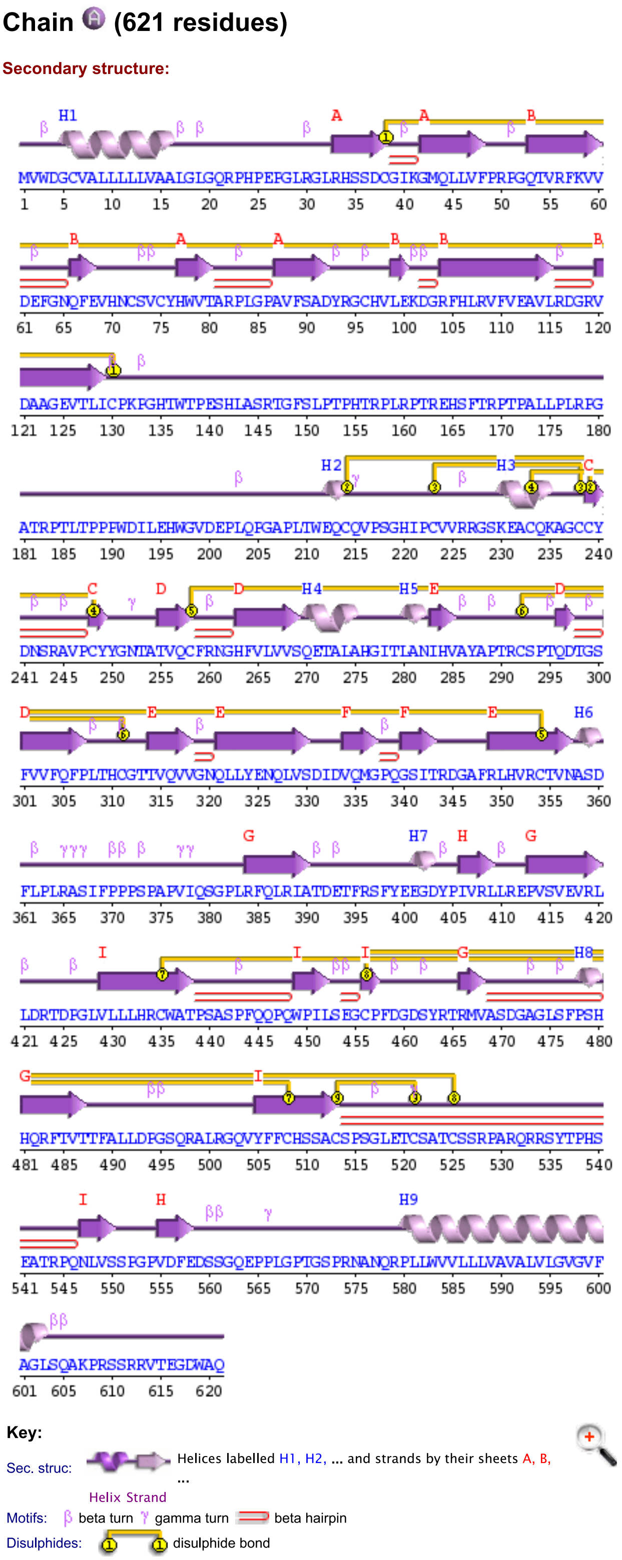


B

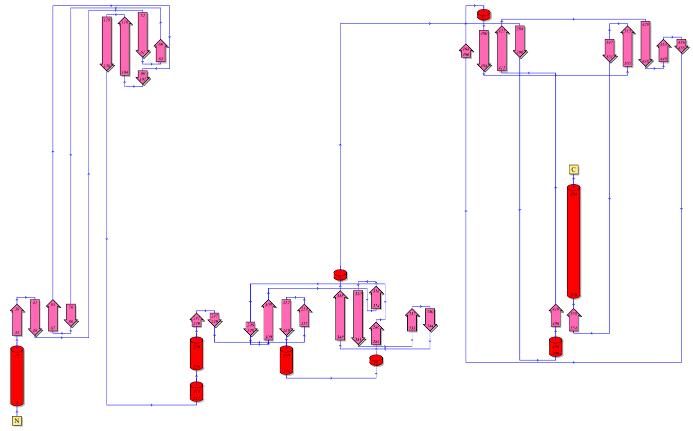

*A. Secondary structure of ZP1 protein*
*B. Topology of ZP1 protein*

### IZUMO1

A

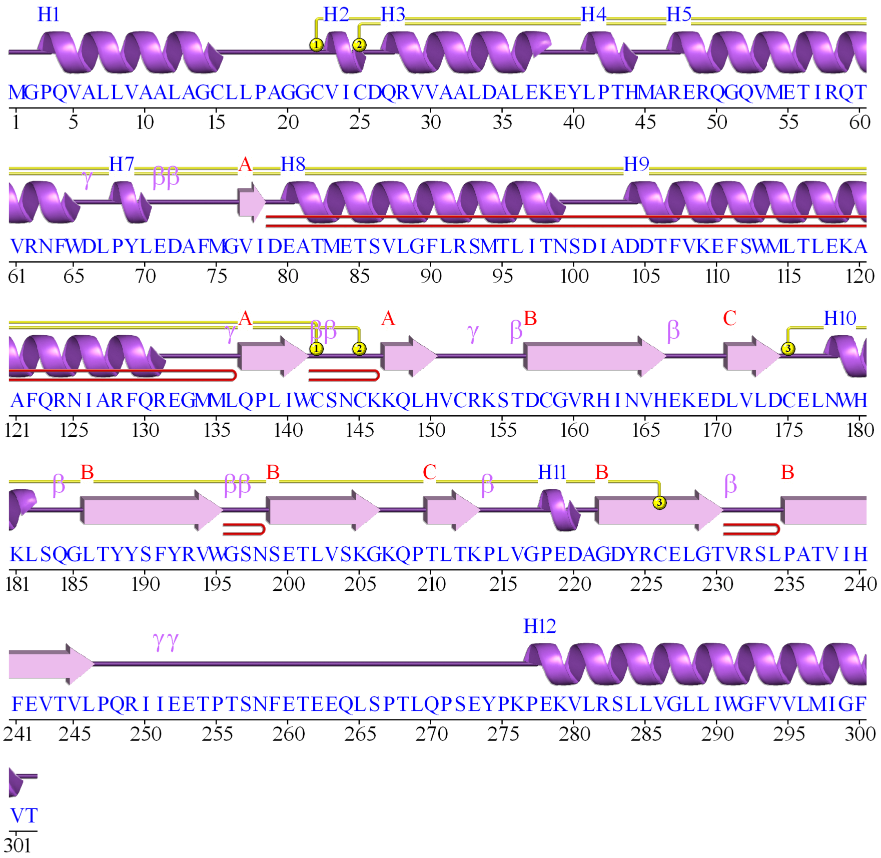


B

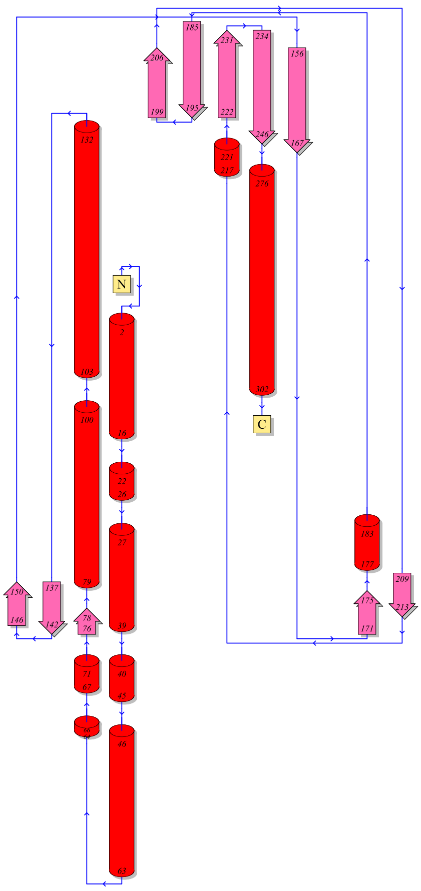


*A. Secondary structure of IZUMO1 protein*
*B. Topology of IZUMO1 protein*

### IZUMO1

A

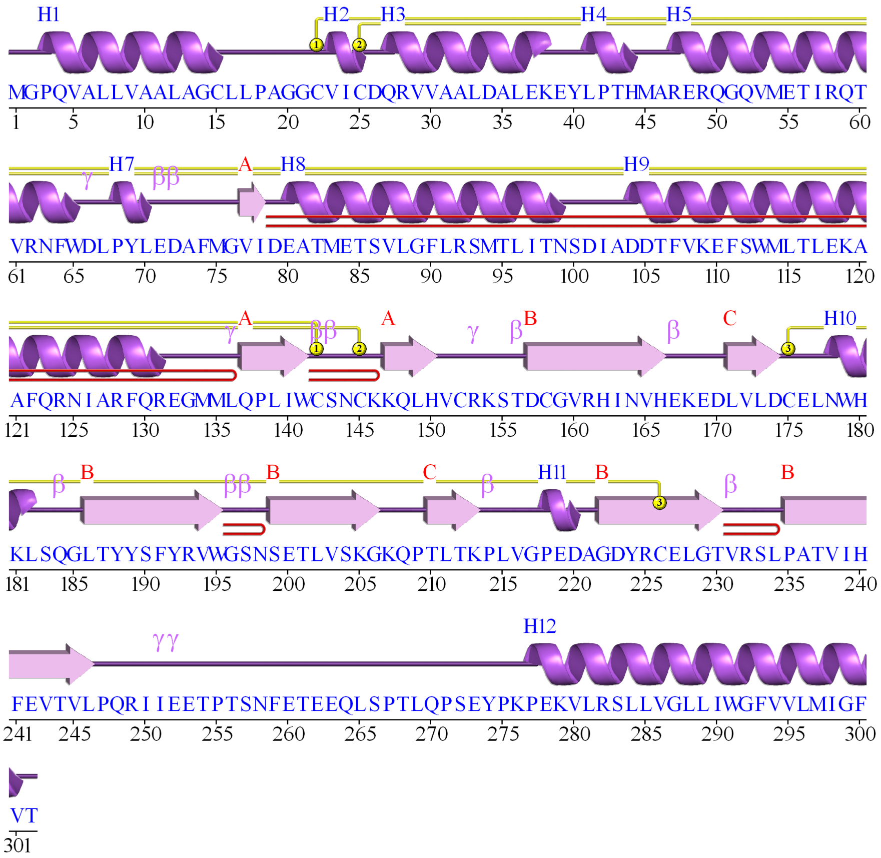


B

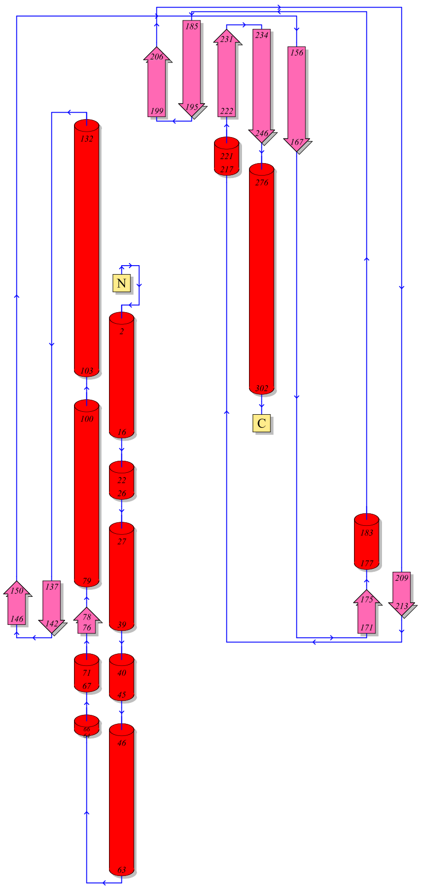


*A. Secondary structure of IZUMO1 protein*
*B. Topology of IZUMO1 protein*

### IZUMO1

*Alignment detail*

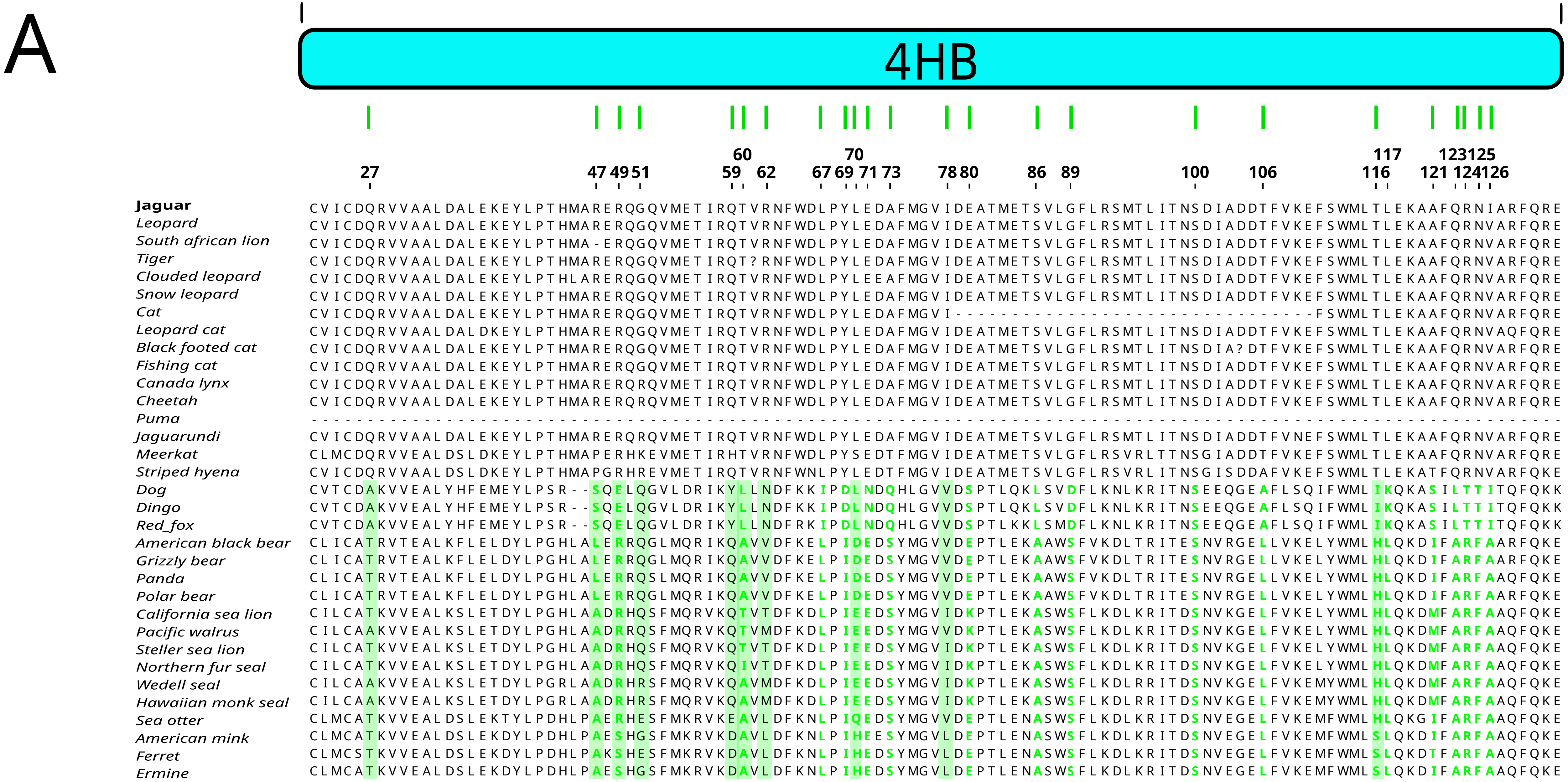


*Alignment detail of the IZUMO1 protein to showcase positively selected sites. Diversifying positive selection in the sub-tree is highlighted by coloring amino acids in bold green lettering, while episodic positive selection on the basal branch of the clade is indicated by a green background.*

### IZUMO1

## B

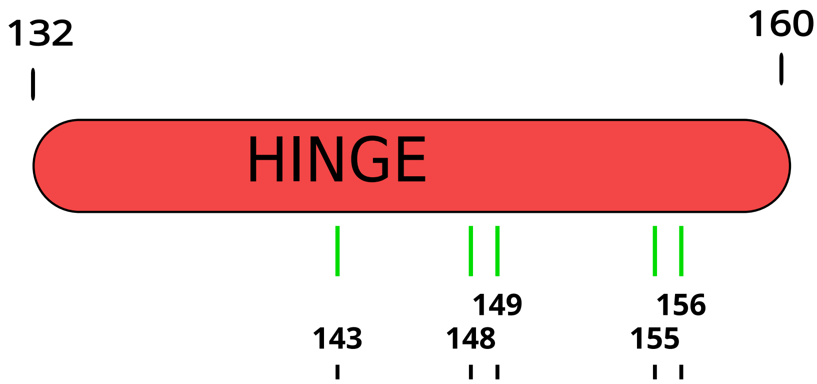


| Species | Alignment (132–160) |
|---|---|
| **Jaguar** | -GMMLQPLIWCSNCKKQLHVCRKSTDCGV |
| *Leopard* | -GMMLQPLIWCSNCKKQLHVCRKSTDCGV |
| *South african lion* | CGMMLQPLIWCSNCKKQLHVCRKSTDCGV |
| *Tiger* | -GMMLQPLIWCSNCKKQLHVCRKSTDCGV |
| *Clouded leopard* | -GMMLQPLIWCSNCKKQLHVCRKSTDCG? |
| *Snow leopard* | -GMMLQPLIWCSNCKKQLHVCRKSTDCGV |
| *Cat* | -GIMLQPLIWCSNCKKQLHVCRKSTDCGV |
| *Leopard cat* | -GIMLQPLIWCSNCKKQLHVCRKSTDCGV |
| *Black footed cat* | -GIMLQPLIWCSNCKKQLHVCRKSTDCG? |
| *Fishing cat* | -GIMLQPLIWCSNCKKQLHVCRKSTDCG? |
| *Canada lynx* | CGMMLQPLIWCSNCKKQLHVCRKSTDCGV |
| *Cheetah* | -GMMLQPLIWCSNCKKQLHVCRKSTDCGV |
| *Puma* | CGMMLQPLIWCSNCKKQLHVCRKSTDCGV |
| *Jaguarundi* | --MMLQPLIWCSNCKKQLHVCRKSTDCGV |
| *Meerkat* | CGTMLQPLIWCSNCKKELHVCRKPYDCGV |
| *Striped hyena* | CGMMLQPLIWCSNCKKQLHVCRKSYDCGA |
| *Dog* | CGQMLQTLIWCHNCLKRIHTCRKSIDCGV |
| *Dingo* | CGQMLQTLIWCHNCLKRIHTCRKSIDCGV |
| *Red_fox* | CGQMLQTLIWCHNCLKRIHTCRKPIDCGV |
| *American black bear* | CGMMLQSLIWCNTCQKQVHTCRKSSDCGV |
| *Grizzly bear* | CGMMLQSLIWCNTCQKQVHTCRKSSDCGV |
| *Panda* | -GTMLQSLIWCNTCQKQVHTCRKSSDCGV |
| *Polar bear* | CGMMLQSLIWCNTCQKQVHTCRKSSDCGV |
| *California sea lion* | CGMMLQSLIWCNTCQKQVHTCRKSSDCGV |
| *Pacific walrus* | CGMMLQSLIWCNTCQKQVHTCRKNSDCGV |
| *Steller sea lion* | CGMMLQSLIWCNTCQKQVHTCRKSSDCGV |
| *Northern fur seal* | CGMMLQSLIWCNTCQKQVHTCRKSSDCGV |
| *Wedell seal* | CGMMLQSLIWCNTCQKQVHTCRKSSDCGV |
| *Hawaiian monk seal* | CGMMLQSLIWCNTCQKQVHTCRKSSDCGV |
| *Sea otter* | CGVMLQSLIWCNSCERQVHACRKTANCGE |
| *American mink* | CGVMLQSLVWCNSCERQVHACRKHPDCGE |
| *Ferret* | CGVMLQSLVWCNSCQRRVHACRKHPNCGE |
| *Ermine* | CGVMLQSLVWCNSCERQVHACRKHPNCGE |

*Alignment detail of the IZUMO1 protein to showcase positively selected sites. Diversifying positive selection in the sub-tree is highlighted by coloring amino acids in bold green lettering, while episodic positive selection on the basal branch of the clade is indicated by a green background.*

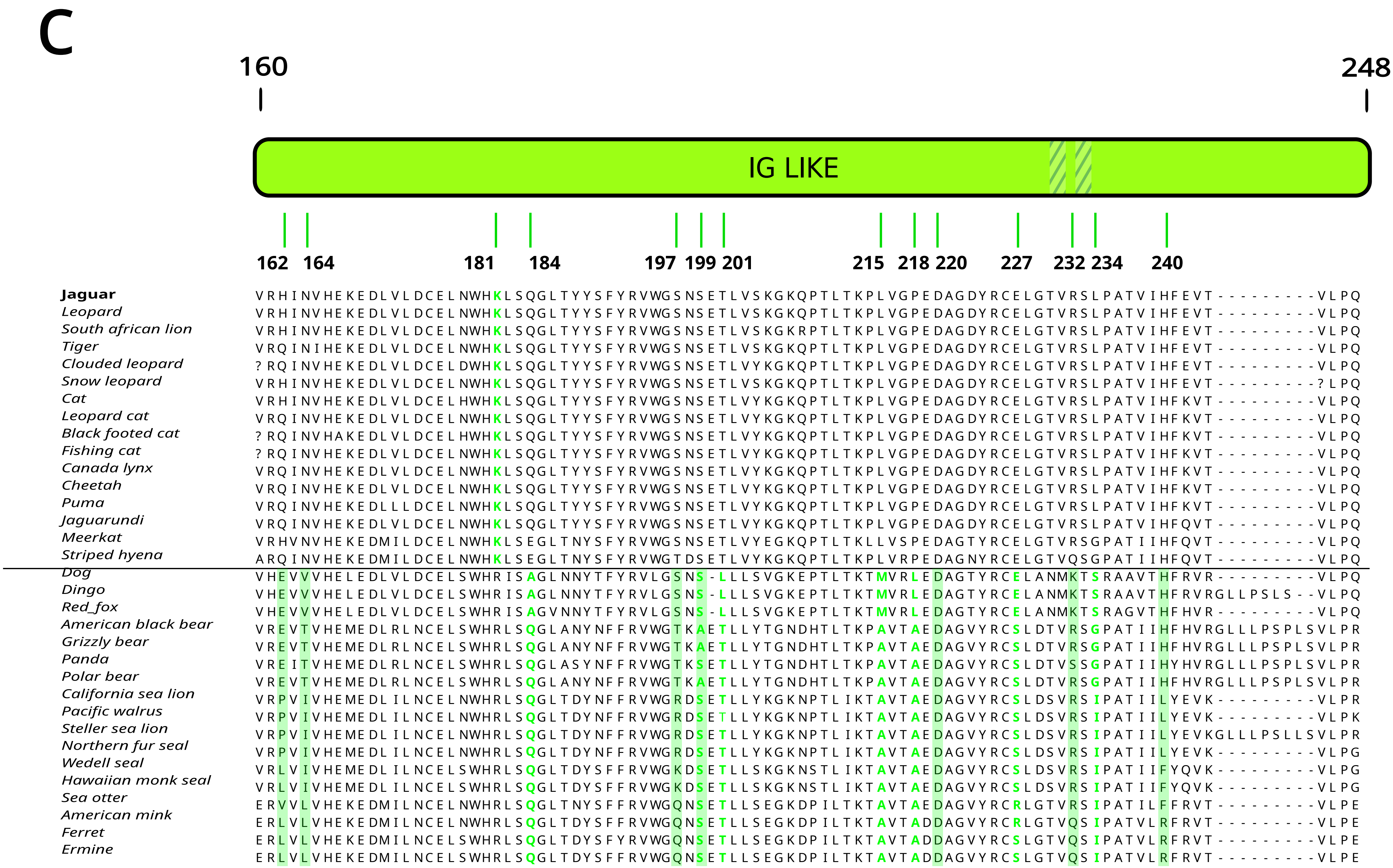


*Alignment detail of the IZUMO1 protein to showcase positively selected sites. Diversifying positive selection in the sub-tree is highlighted by coloring amino acids in bold green lettering, while episodic positive selection on the basal branch of the clade is indicated by a green background.*

### IZUMO1

## D

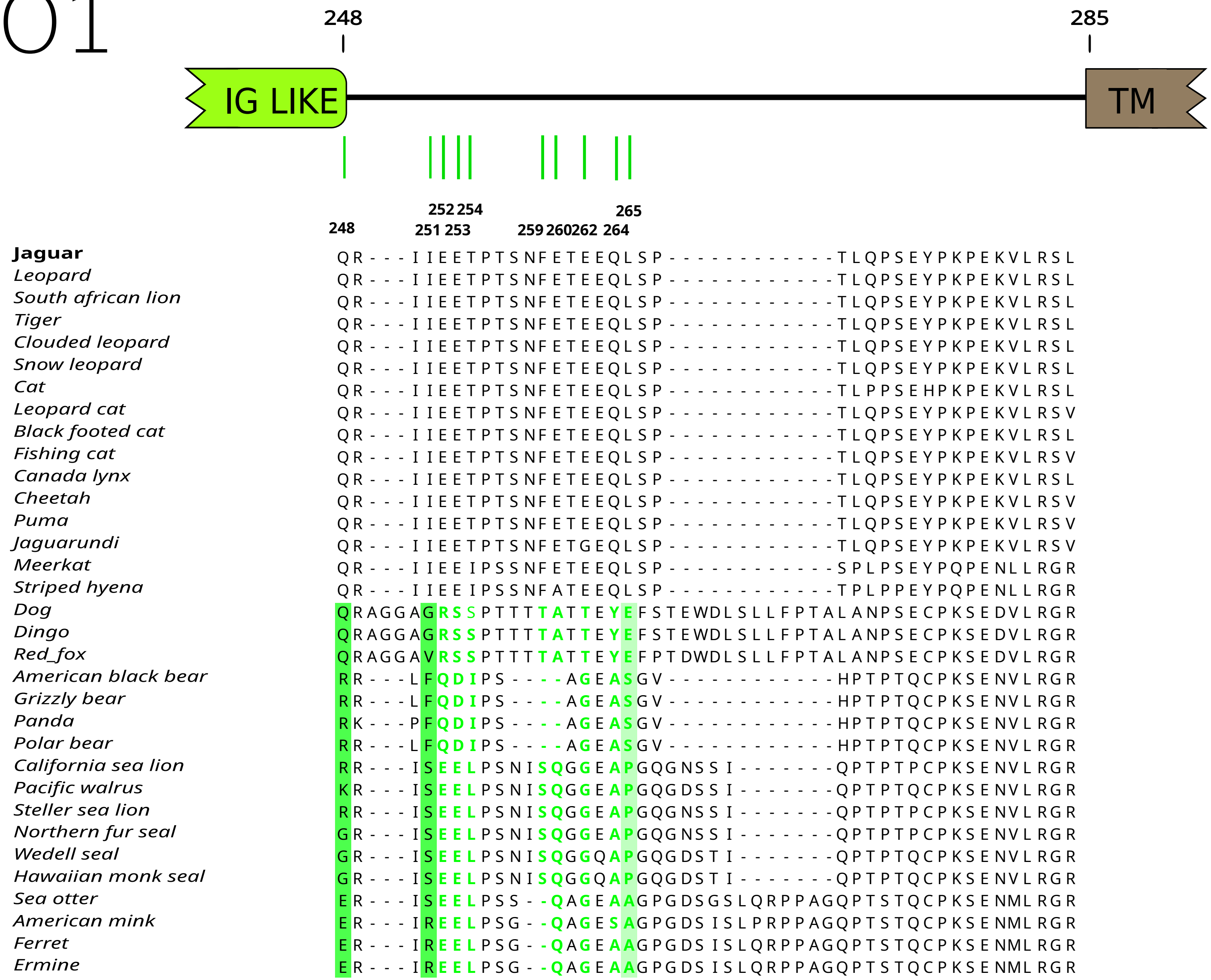


*Alignment detail of the IZUMO1 protein to showcase positively selected sites. Diversifying positive selection in the sub-tree is highlighted by coloring amino acids in bold green lettering, while episodic positive selection on the basal branch of the clade is indicated by a green background.*

### IZUMO1

*Alignment detail*

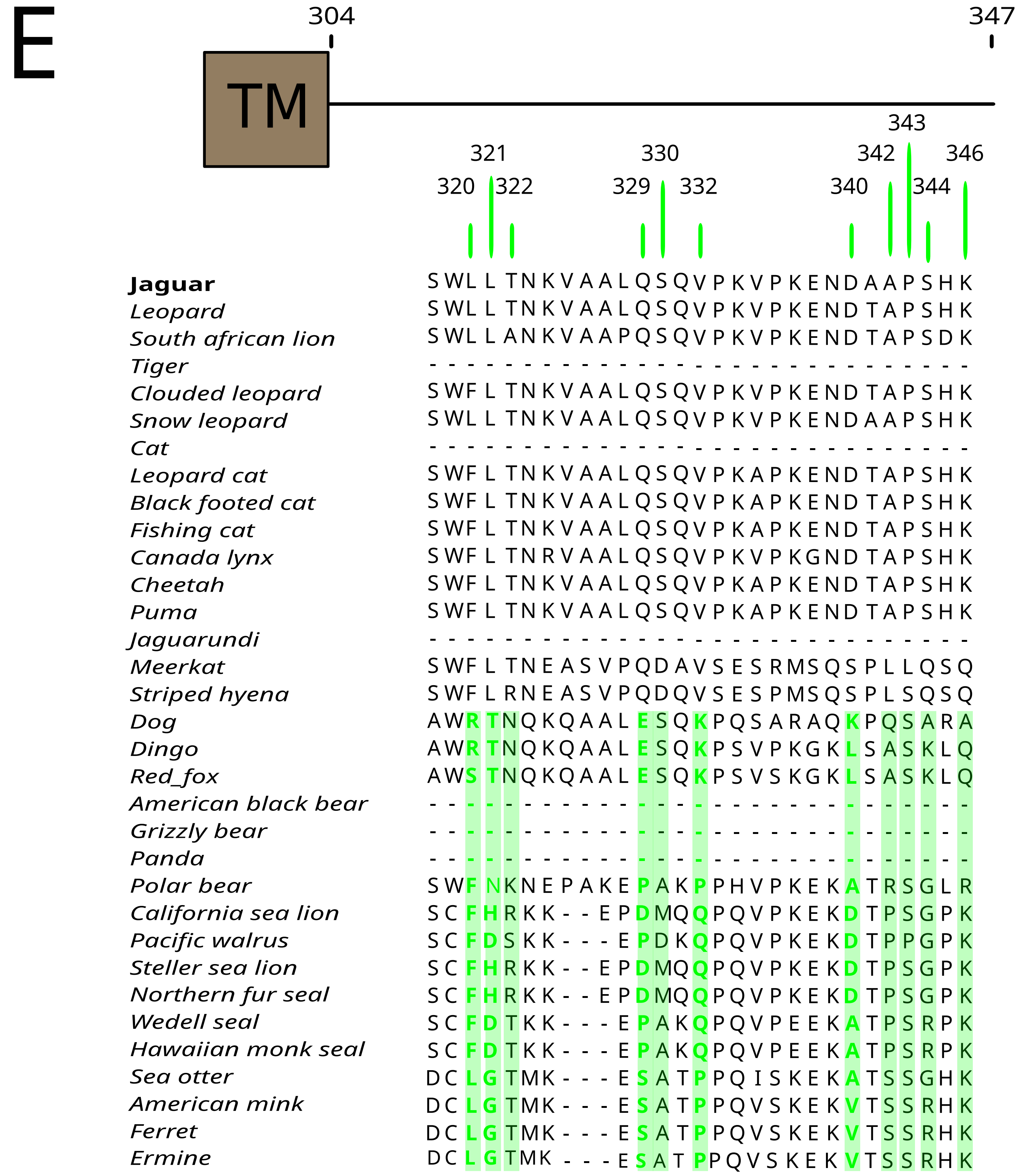


*Alignment detail of the IZUMO1 protein to showcase positively selected sites. Diversifying positive selection in the sub-tree is highlighted by coloring amino acids in bold green lettering, while episodic positive selection on the basal branch of the clade is indicated by a green background.*

**Supplementary table S27: table of positively selected sites within the IZUMO1 protein**

| Protein domain | Position | Identity | | | kind of positive selection | | |
|---|---|---|---|---|---|---|---|
| | | Caniformes | Feliformes | Panterinos | episodic in the basal branch of caniforms | divergent in caniforms subtree | episodic in the basal branch of feliforms |
| 4HB | 27 | A / T | Q | Q | | | |
| | 47 | S / L / A | P / R | P / R | | | |
| | 49 | S/ R / E | R | R | | | |
| | 51 | Q / R / E / G | G / R / K | G / R | | | |
| | 59 | Y / Q / D | Q/ H | Q | | | |
| | 60 | A / T / L | T | T | | | |
| | 62 | L / M / T / V / N | R | R | | | |
| | 67 | P / A | A | A | | | |
| | 69 | K / I / D | Y | Y | | | |
| | 70 | Q / H / E / D / L | L / S | L | | | |
| | 71 | E / N | E | E | | | |
| | 73 | S / Q | A / T | A | | | |
| | 78 | V / L / I | I | I | | | |
| | 80 | E / K / S | E | E | | | |
| | 86 | S / A / L | S | S | | | |
| | 87 | S / A | V | V | | | |
| | 89 | S / D | G | G | | | |
| | 100 | S | S | S | | | |
| | 102 | V / E | I | I | | | |
| | 106 | L / A | T / A | T | | | |
| | 116 | H / S / I | T | T | | | |
| | 117 | L / K | L | L | | | |
| | 121 | I / T / M / S | A / T | A | | | |
| | 123 | A / L | Q | Q | | | |
| | 124 | R / T | R | R | | | |
| | 125 | F / T | N | N | | | |
| | 126 | F / T | V | V / I | | | |
| HINGE | 143 | N / H | S | S | | | |
| | 148 | N / H | S | S | | | |
| | 149 | Q / R | Q / E | Q | | | |
| | 155 | T / H / S / P | S / P | S | | | |
| | 156 | A / P / S / I | T / Y | T | | | |
| IG LIKE | 162 | V / L / P / E | H / Q | H / Q | | | |
| | 164 | L / I / T / V | N | N | | | |
| | 181 | R | K | K | | | |
| | 184 | Q / A | Q / E | Q | | | |
| | 197 | S / T / R / Q / K | S / T | S | | | |
| | 199 | S / A | S | S | | | |
| | 201 | T / L | T | T | | | |
| | 215 | A / M | L | L | | | |
| | 218 | A / L | P | P | | | |
| | 220 | D | D | D | | | |
| | 227 | R / S / E | E | E | | | |
| | 232 | Q / R / S / K | R / Q | R | | | |
| | 234 | I / G / S | L / G | L | | | |
| | 240 | F / R / L / H | H | H | | | |
| | 242 | R / H / E / Q | E / K / Q | E | | | |
| | 248 | E / G / R / Q | Q | Q | | | |
| Interdomain region | 251 | S/ R / F / V / G | I | I | | | |
| | 252 | E / Q / R | E | E | | | |
| | 253 | E / D / S | E | E | | | |
| | 254 | L / I / S | T / I | T | | | |
| | 259 | S / T | F | F | | | |
| | 260 | Q / A | E / A | E | | | |
| | 262 | G / T | E | E | | | |
| | 264 | A / S / IY | Q | Q | | | |
| | 265 | A / P / S / E | L | L | | | |
| | 267 | P / Q / V / S | P | P | | | |
| | 268 | Q / H / L | T / S | T | | | |
| | 270 | L / T / N | Q / P | Q | | | |
| | 273 | E / Q / P | E | E | | | |
| Interdomain region | 320 | F / L / R / S | L / F | L / F | | | |
| | 321 | L / G / D / H / N / T | L | L | | | |
| | 322 | T / R / N | T / A / R | T / A | | | |
| | 329 | S / P / D / E | Q | Q | | | |
| | 330 | A / M / D / S | S / D | S | | | |
| | 332 | P / Q / K | V | V | | | |
| | 340 | A / V / D / L / K | D / S | D | | | |
| | 342 | S / P / R / Q | A / L | A | | | |
| | 343 | S / P | P / S / L | P | | | |
| | 344 | R / G / K / A | G / R / Q / A | S / Q | | | |
| | 346 | K / R / Q / A | K / Q | K | | | |

Table of Sites Detected Under Positive Selection in the IZUMO1 Protein. The table provides the site's position, its location relative to the protein domain, the amino acid identity of the site within the three groups of interest (caniforms, feliforms, and pantherines), and the type of selection (episodic or divergent). Sites highlighted in yellow are those that have been reported to be involved in the interaction with the sperm counterpart, Juno.

### IZUMO1

*Coulombic surface*

ancestral | derived

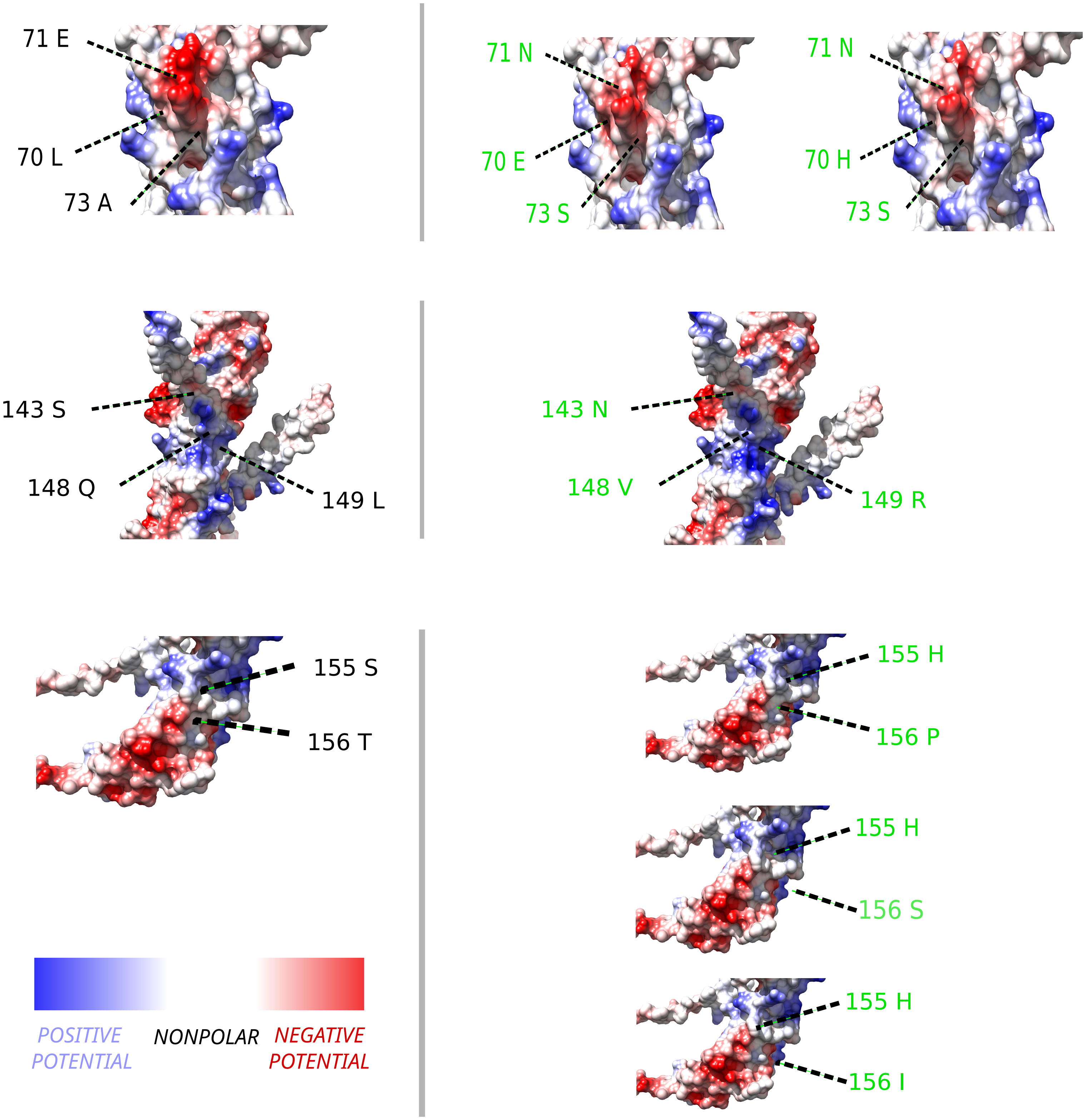


*The ancestral and derived states of positively selected amino acids depicted through Coulombic surfaces. These surfaces illustrate the electrostatic potential as governed by Coulomb's law: red indicates negative potential, blue represents positive potential, and white denotes relatively nonpolar potential.*

### JUNO

A

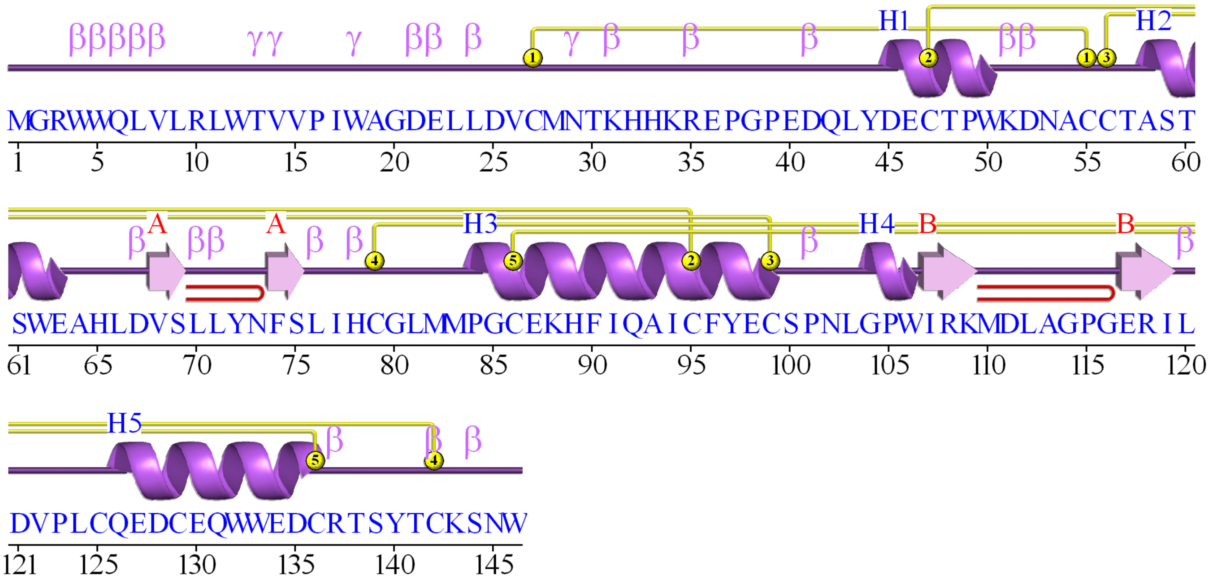


B

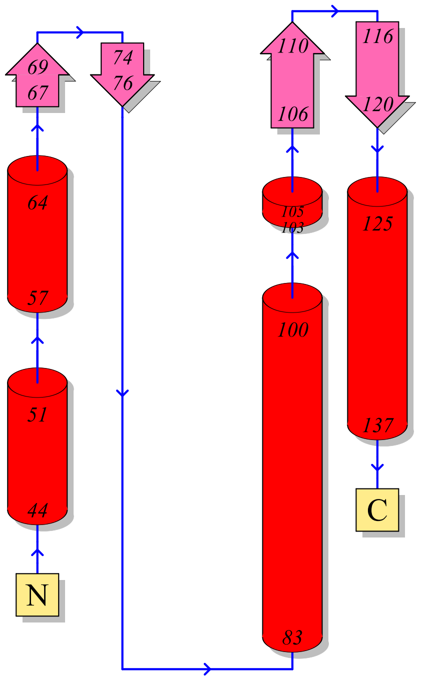


*A. Secondary structure of JUNO protein*
*B. Topology of JUNO protein*

**Supplementary table S31: branch-site positive selection results for control proteins using *Actn1***

| *Actn1 (negative contol)* | | | | | | |
|---|---|---|---|---|---|---|
| | **Log Likelihood** | **LRT** | **p** | **Parameters Estimation** | | |
| **Carnivora** | | | | | | |
| Basal branch | | | | | | |
| Alternative hypothesis | -12050.00624 | | | p0 = 0.99388 | p1 = 0.00612 | p2 = 0.00000 |
| Null hypothesis | -12050.00626 | 0.00003 | 1.000 | ω0 = 0.01547 | ω1 = 1.00000 | ω2 = 1.00000 |
| Subtree analysis | | | | | | |
| Alternative hypothesis | -12049.9043 | | | p0 = 0.99262 | p1 = 0.00598 | p2 = 0.00139 |
| Null hypothesis | -12049.88795 | -0.03271 | 1.000 | ω0 = 0.01525 | ω1 = 1.00000 | ω2 = 1.00000 |
| **Caniformia** | | | | | | |
| Basal branch | | | | | | |
| Alternative hypothesis | -12050.00626 | | | p0 = 0.99388 | p1 = 0.00612 | p2 = 0.00000 |
| Null hypothesis | -12050.00624 | -0.00003 | 1.000 | ω0 = 0.01547 | ω1 = 1.00000 | ω2 = 1.00000 |
| Subtree analysis | | | | | | |
| Alternative hypothesis | -12048.75731 | | | p0 = 0.99061 | p1 = 0.00571 | p2 = 0.00367 |
| Null hypothesis | -12048.74866 | -0.0173 | 1.000 | ω0 = 0.01506 | ω1 = 1.00000 | ω2 = 1.00000 |
| **Feliformia** | | | | | | |
| Basal branch | | | | | | |
| Alternative hypothesis | -12050.00624 | | | p0 = 0.99388 | p1 = 0.00612 | p2 = 0.00000 |
| Null hypothesis | -12050.00624 | 0 | 1.000 | ω0 = 0.01547 | ω1 = 1.00000 | ω2 = 1.00000 |
| Subtree analysis | | | | | | |
| Alternative hypothesis | -12048.8892 | | | p0 = 0.99227 | p1 = 0.00617 | p2 = 0.00156 |
| Null hypothesis | -12049.10985 | 0.4413 | 0.506 | ω0 = 0.01527 | ω1 = 1.00000 | ω2 = 2.14145 |
| **Pantherinae** | | | | | | |
| Basal branch | | | | | | |
| Alternative hypothesis | -12061.17908 | | | p0 = 1.00000 | p1 = 0.00000 | p2 = 0.00000 |
| Null hypothesis | -12050.00625 | -22.34567 | 1.000 | ω0 = 0.01701 | ω1 = 1.00000 | ω2 = 1.00000 |
| Subtree analysis | | | | | | |
| Alternative hypothesis | -12050.00624 | | | p0 = 0.99388 | p1 = 0.00612 | p2 = 0.00000 |
| Null hypothesis | -12050.00624 | 0 | 1.000 | ω0 = 0.01547 | ω1 = 1.00000 | ω2 = 1.00000 |
| ***Panthera onca*** | | | | | | |
| Basal branch | | | | | | |
| Alternative hypothesis | -12050.00799 | | | p0 = 0.99388 | p1 = 0.00612 | p2 = 0.00000 |
| Null hypothesis | -12050.00625 | -0.00348 | 1.000 | ω0 = 0.01546 | ω1 = 1.00000 | ω2 = 1.00000 |

LRT: statistical value of the likelihood ratio test; p0-2 and ω0-2 proportion of sites and dN/dS values estimated for each site class in the selection model (alternative hypothesis). respecively.

**Supplementary table S32: branch-site positive selection results for control proteins using *Tpm3***

| *Tpm3 (positive control)* | | | | | | |
|---|---|---|---|---|---|---|
| | **Log Likelihood** | **LRT** | **p** | **Parameters Estimation** | | |
| ***Panthera onca*** | | | | | | |
| Basal branch | | | | | | |
| Alternative hypothesis | -2577.9999 | | | p0 = 0.89229 | p1 = 0.10658 | p2 = 0.00113 |
| Null hypothesis | -2581.76627 | 7.53274 | 0.00606 | ω0 = 0.01625 | ω1 = 1.00000 | ω2 = 999.00000 |

LRT: statistical value of the likelihood ratio test; p0-2 and ω0-2 proportion of sites and dN/dS values estimated for each site class in the selection model (alternative hypothesis). respecively.
